# Environmental conditions modulate constitutive and induced host immunity contributing to spatial patterns of coral disease susceptibility

**DOI:** 10.1101/2025.11.04.686625

**Authors:** Nicholas P. Jones, Sydney Dutton, Erin N. Shilling, Weston H. Nowlin, Lauren E. Fuess

**Affiliations:** National Coral Reef Institute, Halmos College of Arts and Sciences, Nova Southeastern University, 8000 N Ocean Drive, Dania Beach, Florida, 33004, USA; Texas State University, 601 University Drive, San Marcos, Texas, 78666, USA

**Keywords:** coral reefs, multi-stressor, SCTLD, modeling, innate immunity, climate change

## Abstract

Disease outbreaks have caused mass mortality and threaten the persistence of numerous vulnerable species, including reef-building corals. Spatiotemporal variations in disease prevalence suggest environmental conditions exacerbate disease susceptibility, potentially by compromising host immunity. Understanding how these variations influence disease dynamics and immunity may enable improved prediction of future disease outbreaks. We combined field observations and statistical modeling to identify the environmental drivers of stony coral tissue loss disease (SCTLD) in Florida. We then performed environmental manipulation experiments to confirm the effects of identified factors on immunity in the massive coral *Montastraea cavernosa*. SCTLD susceptibility was influenced most by the interaction between temperature and chlorophyll-a concentration (nutrients proxy) the month prior to disease survey, as well as the interaction between chlorophyll-a and three-month mean PAR. SCTLD prevalence was highest when temperatures were low (<30 °C) and chlorophyll-a concentration exceeded ∼6 mg m^-3^ and/or when PAR and chlorophyll-a concentrations were high. SCTLD severity had a negative relationship with temperature, a colony had a higher probability of dying during the SCTLD outbreak when temperatures stayed below 31.08 °C, a finding in contrast to previous coral disease studies. In the laboratory, coral fragments were exposed to temperature and nutrient manipulation, then immune challenged. Heat stress largely drove suppression of baseline immunity but increased production of some antioxidants, suggesting host stress. Fragments exposed to moderate ammonium concentration induced the strongest immune responses compared to those grown under high or no ammonium conditions. Combined with modeling results, these findings support long-standing hypotheses that coral disease susceptibility is at least, in part, modulated by environmentally induced changes in immunity. Overall, our results enhance the understanding of the impact of environmental conditions on SCTLD outbreaks and on general coral immunity. They also highlight the possibility of increased disease severity as anthropogenic pressure on marine environments increases.

## 1. Introduction

One of the greatest and most universal threats to global biodiversity over recent decades has been outbreaks of epizootic diseases (Daszak et al., 2000). Studies show disease outbreaks have driven significant declines of numerous groups of organisms including amphibians (Fisher & Garner, 2020), echinoderms (Hewson et al., 2014), and bats (Cheng et al., 2021). Notably, these epizootic outbreaks across systems have been increasing in severity and prevalence in recent years (Burge et al., 2014; Burke et al., 2023; Ristaino et al., 2021; Rowley et al., 2024), a pattern that is believed to be driven by the synergistic effects of a rapidly changing environment on hosts and pathogens. Indeed, hotter temperatures (Barris et al., 2018; Burge et al., 2014; Eisenlord et al., 2016; Rodenberg et al., 2024) nutrient enrichment (Klinges et al., 2022; Wear & Thurber, 2015) and other anthropogenically associated stressors (Belasen et al., 2019; Berkhout et al., 2023) have been demonstrated to increase host susceptibility to pathogens, potentially facilitating disease. Furthermore, some of these factors may also impact pathogen virulence (Burge et al., 2014; Cohen et al., 2020; Roussin-Leveillee et al., 2024) contributing to changes in disease. These patterns of association are often visualized in the disease triangle (Scholthof 2007) and have been well documented from an ecological perspective. However, in many systems we lack a fundamental understanding of the cellular and molecular mechanisms which produce these patterns. This lack of fundamental knowledge significantly hinders our ability to accurately predict disease outcomes, manage disease outbreaks, and overall better conserve vulnerable species.

The impacts of disease have been particularly pronounced in marine systems, where numerous keystone species have faced large declines in response to pathogenic outbreaks (Harvell et al., 1999). Amongst the marine species hardest hit are scleractinian corals, which form the basis of coral reef ecosystems (Richardson, 1998). Disease outbreaks have been a primary cause of mass scleractinian mortality on Western Atlantic coral reefs since the 1970s and continue to threaten the persistence of coral populations in this region (Harvell et al., 1999; Randall & van Woesik, 2015; Weil, 2004). While Indo-Pacific coral reefs have not been as badly affected, disease has still been a major source of mortality (e.g., (Page et al., 2023; Sato et al., 2009). Corals are also susceptible to a wide variety of environmental stressors (Maina et al., 2011), many of which also exacerbate disease (Harvell et al., 1999; Vega Thurber et al., 2020), presumably by either compromising the coral host (Lesser et al., 2007; Muller et al., 2008) or increasing pathogen prevalence and virulence (Muller & van Woesik, 2009). Thermal stress (Bruno et al., 2007; Miller & Richardson, 2014; Miller et al., 2009), nutrient enrichment (Vega Thurber et al., 2014) and suspended sediments/turbidity/light (Pollock et al., 2014; Randall & van Woesik, 2015) have been associated with inducing and exacerbating disease outbreaks, likely, at least in part, by compromising coral immunity (Palmer et al., 2010). However, like in many other systems, these relationships have generally been identified through field observations, which identify correlation, but not causation (Aeby et al., 2020; Randazzo-Eisemann et al., 2022; Ruiz-Moreno et al., 2012). Alternatively, some studies have focused on isolated laboratory experiments, which may not reflect ecologically relevant conditions (Palmer, McGinty, et al., 2011; Takagi et al., 2020). Indeed, there is a pressing need to combine both approaches to identify ecologically meaningful environmental variables which may exacerbate disease, experimentally validate their impact on disease susceptibility, and characterize the underlying cellular and molecular pathways driving these patterns. This type of integrative approach will greatly enhance the general understanding of the mechanisms which drive environmental impacts on disease susceptibility and dynamics and improve the modeling and prediction of coral disease outbreaks for conservation applications.

Over two decades of foundational work has significantly enhanced the understanding of coral immune responses, which are now known to be highly homologous to mammalian innate immune systems (Emery et al., 2024). Corals possess a sophisticated repertoire of cellular pattern recognition receptors which aid in pathogen detection and management of microbial symbioses (Emery et al., 2024). These receptors trigger a variety of downstream signaling cascades which result in both cell-mediated and humoral immune responses including phagocytosis, melanization, antimicrobial compound production, and reactive oxygen bursts (and associated host-protective production of antioxidants; (Emery et al., 2024). Many of these responses are known to be impacted by environmental conditions, though there is currently a lack of generalizable rules regarding the relationship between environmental stressors and coral immune performance (Traylor-Knowles & Connelly, 2017). Improved understanding of the link between changing environmental conditions and host coral immune performance is essential for understanding the mechanistic basis of associations between environmental conditions and disease outbreaks in corals.

Disease outbreaks, such as stony coral tissue loss disease (SCTLD), can act as invaluable case studies to investigate the influence of environmental conditions on disease dynamics and host immunity. SCTLD was first reported in 2014 in southeast Florida (Precht et al., 2016) and has since devastated many Western Atlantic coral populations (Alvarez-Filip et al., 2022; Hayes et al., 2022). While the broadscale impacts of SCTLD have been severe, spatiotemporal differences in disease susceptibility and severity suggest local environmental conditions exacerbated SCTLD susceptibility and enhanced coral mortality (Alvarez-Filip et al., 2019; Dahlgren et al., 2021; Williams et al., 2021). To date, no clear environmental pattern has been identified to explain the spatiotemporal variations in SCTLD etiology or epidemiology. Initial outbreak reports suggested an environmental trigger, primarily prolonged heat stress (Jones et al., 2021; Precht et al., 2016), but there are suggestions that the subsequent spread and peaks in SCTLD elsewhere in Florida were not associated with heating (Muller et al., 2020; Williams et al., 2021). Furthermore, elsewhere in the Caribbean, poor water quality has been implicated as exacerbating coral mortality rates (Alvarez-Filip et al., 2019; Alvarez-Filip et al., 2022). Regardless of the exact trigger, it remains plausible that environmental conditions predisposed coral communities to disease, perhaps through impacts on coral immunity, as has been found in other diseases and in other locations (Ban et al., 2014; Lapointe et al., 2019; Lesser et al., 2007; Muller et al., 2018; Muller & van Woesik, 2009; van Woesik & Randall, 2017; Voss & Richardson, 2006). Furthermore, it is highly likely that environmental perturbations will continue to increase disease susceptibility and reduce coral recovery potential (Jones & Gilliam, 2024)

Here, we used the SCTLD outbreak as a case study to apply an integrative modeling and experimental approach to identify and validate environmental conditions which might exacerbate coral disease in the massive Caribbean coral, *Montastraea cavernosa*. As a moderately susceptible species to SCTLD with high variability in both temporal (Shilling et al., 2021) and spatial (Aeby et al., 2019) patterns of SCTLD outbreaks, *M. cavernosa* was an ideal species for such a study. We applied thorough modeling approaches to first identify factors which predicted disease susceptibility and severity across *M. cavernosa’s* range in Florida. We then experimentally validated the impacts of the identified factors on constitutive (baseline) and induced immune responses of the coral host *ex situ*. Combined, our results identify environmental factors which have significant impacts on coral disease susceptibility and demonstrate mechanistically that these associations are likely linked to changes in both constitutive and induced immune responses. These findings are a significant step forward in advancing the knowledge needed to predict coral futures under rapidly changing environmental conditions and increasing disease prevalence.

## 2. Materials and Methods

### 2.1. Modeling of disease susceptibility and severity

Disease susceptibility was quantified as SCTLD prevalence (the proportion of *M. cavernosa* colonies with SCTLD) at individual sites surveyed as part of the Coral Reef Evaluation and Monitoring Project (CREMP), Southeast Florida Coral Reef Evaluation and Monitoring Project (SECREMP), and the Florida Reef Resilience Program’s (FRRP) Disturbance Response Monitoring (DRM). If sites had multiple transects surveyed, *M. cavernosa* abundance and SCTLD prevalence were summed to the site level. As the spread of SCTLD varied spatiotemporally across Florida’s Coral Reef (FCR), prevalence data was filtered to capture the peak SCTLD outbreak period in each FCR subregion using previously documented timelines (Hayes et al., 2022; Muller et al., 2020; Walton et al., 2018; Williams et al., 2021) and by visually assessing spatiotemporal variation in SCTLD prevalence. The peak SCTLD outbreak period was considered to be from 2014-2016 throughout the Kristin Jacobs Coral Aquatic Preserve (Coral AP): Martin, Palm Beach, Deerfield, Broward, Miami, and Biscayne subregions; from 2016 to 2018 in the Upper and Middle Keys, from 2018 to 2019 in the Lower Keys and Marquesas; from 2019 to 2022 in the Marquesas-Tortugas transition zone and from 2020 to 2022 in the Dry Tortugas (Figure 1a).

**Figure 1.**
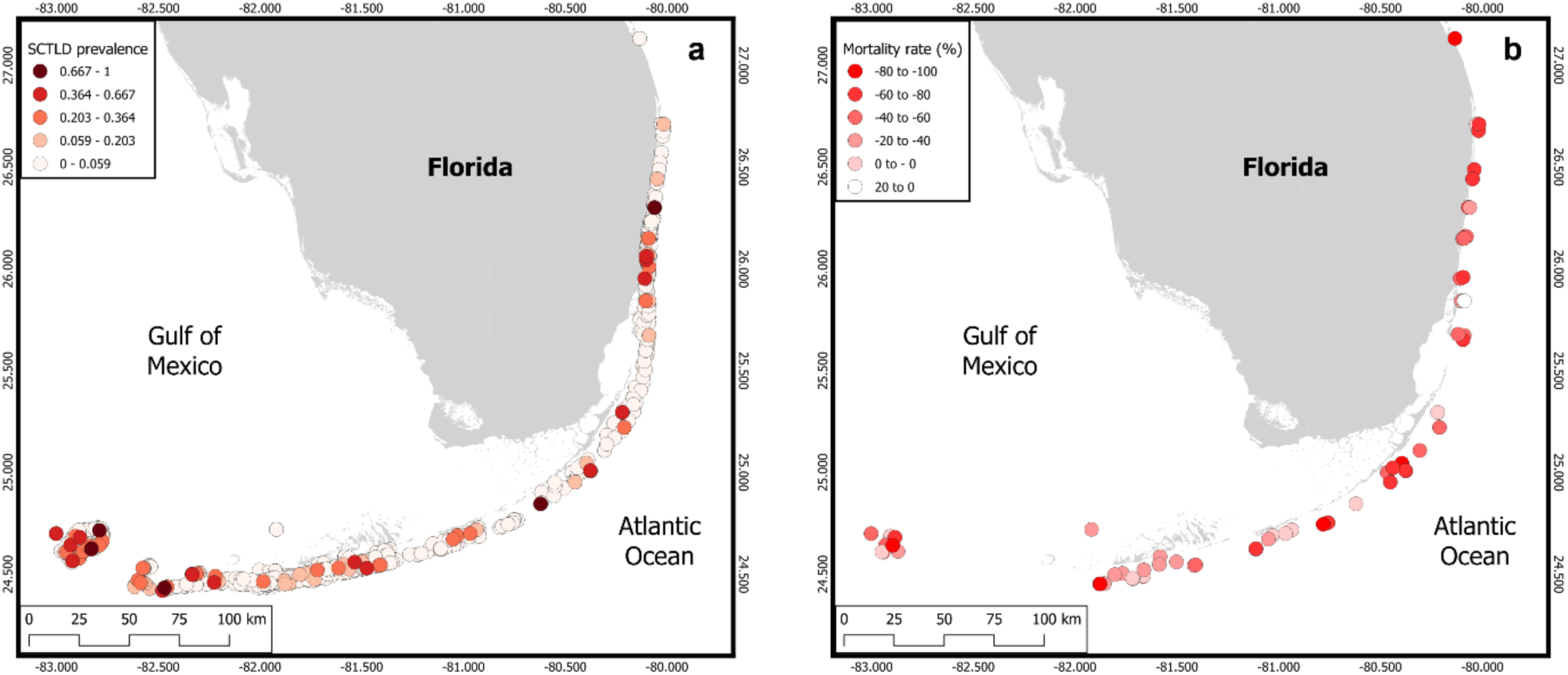
a) SCTLD susceptibility (as prevalence) and b) SCTLD severity (mortality rate) across Florida’s Coral Reef. SCTLD susceptibility was calculated as the proportion of *Montastraea cavernosa* colonies recorded with SCTLD during surveys conducted during the peak outbreak years in each subregion. 9846 colonies were surveyed to quantify disease susceptibility. SCTLD severity was calculated as the two-year relative change in *M. cavernosa* colonies at CREMP or SECREMP sites following the peak in SCTLD prevalence. Individual points are survey sites where darker colors indicate higher SCTLD prevalence or mortality rate. Negative mortality rates indicate a decline in *M. cavernosa* abundance, with -100% indicating all colonies at a site died. Figure created in QGIS v3.40 (Team, 2025).

Disease severity was quantified as the mortality rate (the relative change in *M. cavernosa* abundance) during the initial two years of the SCTLD outbreak at CREMP and SECREMP sites. Abundance at CREMP sites was counted on four, 10 m x 1 m permanent transects, and at SECREMP sites on four, 22 m x 1 m permanent transects. At each site, *M. cavernosa* abundance was summed per site per year. The mortality rate was calculated as the change in abundance from year 0 to year 2, divided by the initial abundance (i.e., relative change (% yr^-1^)). Only colonies ≥4 cm maximum diameter were counted to avoid capturing colonies that recruited during the disease outbreak. Two years was considered sufficient time for colonies which developed SCTLD during the peak outbreak period to have died while simultaneously avoiding capturing recruits that grew into the adult dataset by the second year (Jones & Gilliam, 2024). To capture the peak in mortality rate at each site, mortality rates were calculated for multiple two-year periods and the period with the largest relative change in *M. cavernosa* abundance chosen for modeling. The period generally stayed consistent within a subregion such that the chosen period was 2014 to 2016 in Broward, Miami and Biscayne, 2015 to 2017 in Martin, Palm Beach and Deerfield, 2016 to 2018 in the Upper and Middle Keys, 2017 to 2019 in the Lower Keys and 2020 to 2022 in the Dry Tortugas (Figure 1b).

### 2.2 Quantifying environmental predictors

Environmental data prior to and during the peak SCTLD outbreak were compiled from in situ and satellite data. Specific environmental predictors which have previously been identified to affect stress response in stony corals (Caldwell et al., 2024; van Woesik & Randall, 2017), were then calculated at each benthic sampling site. Data from NASA’s Moderate Resolution Imaging Spectroradiometer satellite (MODIS Aqua) was used to measure solar irradiance, diffuse attenuation coefficient (turbidity proxy, as used in (Caldwell et al., 2024), and chlorophyll-a concentration (nutrients proxy, as used in (Muller et al., 2020). Irradiance was obtained as the surface downwelling photosynthetic flux in air (Photosynthetically Available Radiation (PAR) einstein m^-2^ s^-1^) and diffuse attenuation coefficient as kD490 (m^-1^). Data were extracted from NASA’s Ocean color database (https://oceancolor.gsfc.nasa.gov/l3/) as the monthly mean for each metric at 1/25^th^ resolution throughout Florida. Temperature data was obtained from satellite data and *in situ* sampling. Degree heating week data was obtained from NOAA’s Coral Reef Watch (Watch, 2019). Daily in situ temperature data was collected by HOBO v2 temperature loggers at CREMP and SECREMP sites throughout FCR.

The environmental regime at each benthic sampling site was calculated using a spatial join with the in situ and satellite data. Each benthic site was joined with the three closest satellite sites and the monthly mean of each variable calculated per benthic site. To calculate the temperature regime at DRM sites which were used in the disease susceptibility analysis, a spatial join was made to the nearest CREMP/SECREMP site. If the closest CREMP/SECREMP site had missing data, then a spatial join was made to the closest two sites. After the database with the environmental regime at each benthic site was created, multiple environmental predictors were created to represent conditions either leading up to the disease susceptibility surveys or in the time between disease severity surveys.

For each environmental variable used in the disease susceptibility analysis (chlorophyll-a concentration, PAR, kD490 and temperature) the maximum and mean values were quantified within the month prior, within the prior three months and within the prior year to the benthic survey. To further capture the temperature regime at a site, the minimum temperature over one month, three months and one year prior to the benthic survey and specific thermal stress predictors were calculated. Thermal thresholds were calculated independently for each subregion using modelled SST data from the Hybrid Coordinate Ocean Model (HYCOM) from 2014 to 2022, as 1 °C above the maximum of the mean summertime (July-September) SST, or 1 °C below the minimum of the mean wintertime (January-March) SST as per Jones et al. (Jones et al., 2020). The heat stress or cold stress durations were calculated for each site as the number of days in situ water temperature exceeded thermal thresholds in the month, three months and year before a benthic survey. This gave 31 environmental predictors, which were tested for collinearity by calculating the Spearman’s rank correlation coefficient, which is well suited to assess non-linear relationships and data which are non-normal. Correlations were considered significant at a threshold of 0.8; in cases of collinearity one of the predictors was removed prior to statistical analysis. An attempt was made to keep at least one predictor of a variable at each time point. All chlorophyll-a and kD490 predictors were highly collinear and all removed except maximum chlorophyll-a the month prior to the survey, as this was hypothesized to have the greatest effect on disease susceptibility. Phytoplankton, whose preferred nitrogen source is ammonium (Dortch, 1990), are substantial contributors to both measurements in coastal waters (Fraser, 1998). It is likely that either would have acted as a suitable nutrient proxy, but as chlorophyll-a is more frequently used (e.g., (Muller et al., 2020) it was retained. The removal of other collinear predictors left 15 environmental predictors which were used in statistical analysis (Table 1).

**Table 1.** Environmental summary of predictors used in disease susceptibility full random forest model. Variable = type of environmental variable, predictor = metric of that variable, time period = temporal duration predictor calculated over. e.g., chlorophyll-a maximum, is the maximum chlorophyll-a concentration within one month prior to the disease survey.

| Variable | Predictor | Time period | Mean | SD |
| --- | --- | --- | --- | --- |
| Chlorophyll-a ($\text{mg m}^{-3}$) | Maximum | One month | 3.82 | 5.12 |
| PAR ($\text{Einstein m}^{-2} \text{ day}^{-1}$) | Maximum | One month | 48.16 | 6.11 |
| PAR ($\text{Einstein m}^{-2} \text{ day}^{-1}$) | Maximum | Three months | 56.68 | 2.50 |
| PAR ($\text{Einstein m}^{-2} \text{ day}^{-1}$) | Maximum | One year | 58.23 | 1.93 |
| PAR ($\text{Einstein m}^{-2} \text{ day}^{-1}$) | Mean | Three months | 51.74 | 3.16 |
| PAR ($\text{Einstein m}^{-2} \text{ day}^{-1}$) | Mean | One year | 43.42 | 1.52 |
| Temperature ($^{\circ}\text{C}$) | Maximum | One month | 30.75 | 1.01 |
| Temperature ($^{\circ}\text{C}$) | Maximum | One year | 31.36 | 0.61 |
| Temperature ($^{\circ}\text{C}$) | Mean | Three months | 29.44 | 1.30 |
| Temperature ($^{\circ}\text{C}$) | Mean | One year | 27.13 | 0.61 |
| Temperature ($^{\circ}\text{C}$) | Minimum | One year | 21.51 | 1.70 |
| Thermal stress (days) | Heat stress | One month | 0.93 | 2.77 |
| Thermal stress (days) | Heat stress | Three months | 2.44 | 5.89 |
| Thermal stress (days) | Cold stress | One year | 1.99 | 5.43 |
| Thermal stress (days) | Degree Heating Weeks | At survey | 1.34 | 1.81 |

For the disease severity analysis, the same types of environmental variables were used, but environmental predictors (e.g., maximum or mean temperature) were calculated over the year leading up to the initial benthic survey and in the two years between survey events. This gave 23 environmental predictors, which were tested for collinearity and removed in the same way as described above. The removal of collinear predictors gave 14 environmental predictors which were used in statistical analysis (Table S1).

### 2.3 Statistical analysis: modeling environmental drivers of disease

To assess the relationship between disease susceptibility and disease severity (proportion of colonies that died during the SCTLD outbreak) with environmental predictors, univariate modeling was conducted in a two-step process in R 4.4.1 (R Core Team, 2024). First, for each response metric, a random forest regression model was used to identify the most important predictors of variation in each response variable using the randomForest function from the randomForest package (Liaw & Wiener, 2002). The predictors which accounted for 75% of the variation in the full random forest model were retained and a backwards stepwise regression was performed to identify the most important environmental predictors for stage two using the R^2^. Partial regression plots were inspected for any evidence of quadratic relationships. The selected predictors from the random forest were then modelled as either a binomial Generalized Linear Model (GLM), or binomial Generalized Linear Mixed Model (GLMM), which incorporated any potentially meaningful interactions between predictors.

Disease susceptibility (n = 939 sites) was modelled as SCTLD prevalence (the proportion of diseased *M. cavernosa* colonies at the time of sampling) in both the random forest and GLMM. In the GLMM (created using the package glmmTMB (Brooks et al., 2017), the abundance of *M. cavernosa* at a site was fitted as weights. Sub-region nested within year was fitted as a random effect to account for spatiotemporal variation in disease spread, as sites close together were predicted to be more likely to encounter SCTLD at the same time. Disease severity (n = 62 sites) was modelled as mortality rate (the proportion of *M. cavernosa* colonies which died over the two-year peak outbreak period). In the random forest regression model, the response variable was fitted as the relative change in abundance (% yr^-1^). In the GLM, the response variable was fitted as the proportion that died (i.e., change /initial abundance), with the weights as the initial *M. cavernosa* abundance at the site. Model selection was conducted using a backwards stepwise approach from the full model, containing all predictors and meaningful interactions, by inspecting model summaries and comparing the Akaike Information Criteria (AIC).

Random forest model reliability and goodness of fit were visually assessed by plotting the cross-validation error rate against the number of trees, the root mean squared error (calculated using the out of bag samples) against fitted predictions and fitted model predictions against the observed data. The minimum adequate GLM was validated by plotting deviance residuals against fitted values, and deviance residuals against each significant variable in the fitted model. The fitted GLMM was validated using the package DHARMa with residual diagnostics, including overdispersion, normality and homogeneity of variance, conducted on the fitted model (Hartig, 2017). Overdispersion was detected in the disease severity GLM, the model refit with a quasibinomial distribution and model selection repeated as above. Two outliers were detected and removed. Model validation indicated no further problems. The variance inflation factor (VIF) was used to check fitted models for multicollinearity and found low correlation for the main predictors (VIF < 4) indicating no problems.

### 2.4 Coral collection and husbandry

Fragments of nine unique genets of *M. cavernosa* were collected in Fort Lauderdale, Florida and shipped to Texas State University in early February 2025. Due to shipping delays, corals arrived in various health states and were immediately screened upon arrival for abrasions, lesions, necrosis, and mortality. Fragments with mortality greater than 50% were removed from experimental groups. Remaining fragments were then tagged based on genotype and placed in recirculating tanks with artificial sea water (ASW) made using Instant Ocean Reef Crystals for long term husbandry and experimentation. Prior to experimentation, corals were maintained in this system at conditions mirroring the origin reef environment: 35 ppt salinity, 24°C, 7.2 dKH, 400 ppm Ca, 1400 ppm Mg, and 0 ppm NH_4_, NO^3^, and NO_2_. Light levels were kept at approximately 50 μM/m^2^/s for 11.5 hours daily with a 30-minute ramp up/down period each morning and evening synced to mimic sunrise and sunset in Fort Lauderdale. Corals were maintained in these conditions for an acclimatization period of two and a half weeks. During the first week of acclimatization, corals were dosed with RESTOR (Brightwell Aquatics) and experienced daily 50-75% water changes to account for shipping stress. Following this period, corals were fed twice weekly with Artemia shrimp, followed by 20% water changes. These conditions were maintained through the duration of the experiment.

Two weeks after arrival, corals were slowly acclimatized to average summer temperatures on the FCR. This involved a gradual temperature increase (1 °C every two days) from temperature at collection, ∼24 °C, to a final temperature of 28°C. The control temperature of 28 °C was chosen based on modeling results. A clear shift in the relationship between chlorophyll concentration (nutrients proxy) and disease susceptibility was observed at 30 °C; consequently control (28 °C) and heat stress (32 °C) temperatures equidistant from that shift point were chosen for the experimental portion. During this transition, approximately two and a half weeks after arrival, corals were fragged using a Gryphon Diamond Band Saw. All received fragments were cut into approximately 3 x 3 cm fragments and tagged based on original genotype. Corals were then allowed to recover from fragmentation for two additional weeks. Experimentation was started approximately one week following the end of the ramp to summer temperatures.

### 2.5 Experimental design: Environmental manipulation

After a total acclimatization period of roughly one month, coral fragments were divided into groups to test the impacts of variation in temperature and nutrients (factors identified via modelling as having the strongest impacts on disease susceptibility and severity) on coral immune responses. Two coral fragments of each genotype were randomly assigned to one of six tanks representing a full factorial combination of the two factors: temperature (ambient or heat) and nutrient enrichment (none, mid, high; Figure 2; Table S2). Tanks were isolated from the recirculating system for the duration of the experiment to prevent cross-contamination of temperature or ammonium treatment. Individual heaters were used to raise temperatures in the three heat stress tanks; temperature was ramped from 28 °C to 32 °C at a rate of 1 °C per day and then held at 32 °C for the duration of the experimental period (∼1 month). Temperature in all tanks was monitored throughout the duration of the experiment via HOBO Onset Data Loggers.

**Figure 2.**
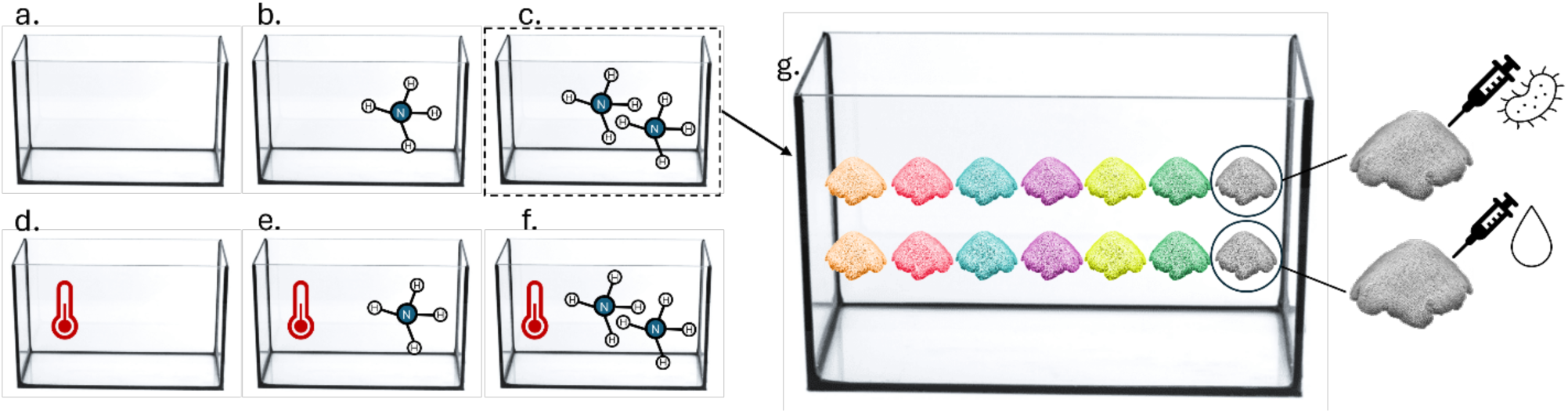
Experimental set up showing environmental manipulations: a. full control (28°C and 0.0 mg/L NH4), b. ambient temperature (28°C) and mid-nutrients (0.01 mg/L NH4), c. ambient temperature and high nutrients (0.05 mg/L NH4), d. heat only (32°C), e. heat and mid nutrients, f. ambient temperature and high nutrients, g. representative tank showing the fourteen frags (two from each genet) in each tank, and how they were split at the end of the environmental manipulation for the immune challenge.

Nutrient enrichment was conducted via the addition of ammonium with concentrations selected to reflect environmental conditions on FCR (Jones & Gilliam, 2024). Ammonium was chosen to represent chlorophyll-a as it is the preferred nutrient source for zooxanthellae and corals (Carmignani et al., 2025; Morris et al., 2019), and has been used as a proxy for nutrients in similar previous experiments (Fuess et al., 2020). At the start of the experiment, nutrient amended tanks were dosed with ammonium chloride (NH_4_Cl, CAS # 12125-02-9) to reach their respective dosage. After initial dosing, levels were measured twice a week in all six tanks using a modification of the phenate method (Solórzano, 1969). The modification involved using sodium citrate (Na_3_C_6_H_5_O_7_, CAS # 6132-04-3) as a complexing agent to eliminate interference of calcium and magnesium (APHA 2023). After each bi-weekly testing of ammonium concentrations, tanks were dosed as needed to maintain the desired concentration. Additionally, when water changes were conducted on feeding days, the ASW used to replace the water removed was dosed with NH_4_Cl to match the target concentration for the respective tank (i.e. the mid-nutrient tank was given ASW at a concentration of 0.01 mg/L of NH_4_). Temperature and nutrient enrichment treatments were maintained for one month before immune challenge experiments to mimic disease susceptibility modeling results.

### 2.6 Experimental design: Immune challenge

Following the one month of environmental manipulation, we conducted an experimental immune challenge. Coral fragments within each treatment tank were randomly assigned to one of two treatments: placebo or immune stimuli, allowing a full factorial combination of treatments (temperature x nutrient x immune challenge) with one fragment per genet in each of the 12 treatment groups. Each coral fragment was placed into an aerated, individual 500 mL, autoclaved plastic beaker with ∼400 mL of 0.2 μm filtered ASW at 35 ppt. For ease of experimentation, ammonium treatment was not continued through the immune challenge portion of the experiment. Individual beakers were randomly placed into larger aquaria by heat treatment, which served as water baths maintaining a temperature of 28°C or 32°C (respective of their original treatment) within the beakers. Coral fragments were allowed to acclimate in the beakers for two hours and were then injected with 500 uL of either the immune stimuli or placebo treatment: heat-killed bacteria, *Vibrio coralliilyticus* at 1 x 10^8^ CFU/mL in sterile ASW, or sterile ASW (vehicle control/placebo). *Vibrio coralliilyticus* is a known coral pathogen commonly used in experiments investigating immune response (Santos Ede et al., 2011; Takagi et al., 2020). Injections were split across 5 locations in the fragment with ∼100 uL injected in each location, for a total injection volume of 500 uL. Corals were then returned to their containers and incubated for 12 hours, at the end of which samples were flash frozen for downstream analyses.

### 2.7 Sample processing and analysis

Host enriched protein extracts were collected from each sampled coral following established protocols (Changsut et al., 2022; Fuess et al., 2016). Briefly, tissue was removed from frozen coral fragments using a Paasche airbrush and 100 mM Tris + 0.05mM DTT (pH 7.8) buffer. The resultant slurry was processed to generate three unique aliquots of tissue for the following analyses: 1) symbiont density estimation (slurry removed from a fixed surface area of the sample), 2) melanin concentration assessment, and 3) protein activity assays. Symbiont density aliquots were washed to remove cell debris and then resuspended in deionized water and stored at 4°C until counts were performed. The melanin aliquot was placed into a pre-weighed 1.5 mL tube, flash frozen, and stored at -20° C until analysis. Finally, the protein activity aliquot was split across two tubes and stored at -80 °C until analysis.

Symbiont density was estimated in each sample via counting in triplicate using a standard hemocytometer. Brightfield and chlorophyll-a fluorescence images were captured on a Biotek Cytation 1 and counts were estimated using ImageJ (Schneider et al., 2012). Counts were averaged within a sample and standardized to tissue area.

Immune assays were conducted following established procedures (Changsut et al., 2022; Fuess et al., 2016). For each assay, samples were run in triplicate and outputs were measured on the BioTek Cytation 1 imaging reader with appropriate negative controls, unless specified otherwise. Prior to conducting immunological assays, protein concentration was measured for each using a Red660 assay (Changsut et al., 2022; Fuess et al., 2016). Measurements from subsequent assays were all standardized by protein concentrations unless otherwise indicated.

Three components of the coral immune response were measured: antioxidant activity, melanin synthesis, and antibacterial compound activity. The coral immune response results in the production of harmful reactive oxygen species (ROS) that can damage both pathogens and host coral. Consequently, antioxidants are often produced by host corals as part of the immune response to prevent immunopathology (Emery et al., 2024). Antioxidants may also be produced to combat ROS produced during periods of thermal or other types of stress (Gardner et al., 2017; Morris et al., 2019). Two types of antioxidant activity were measured following standard protocols: catalase and peroxidase (Changsut et al., 2022; Fuess et al., 2016). Corals, like many other invertebrates, employ melanin synthesis as a direct defense against pathogens. Melanin may have direct cytotoxic effects on pathogens or may be used to encapsulate pathogens and heal wounds (Emery et al., 2024). Melanin synthesis is the result of the prophenoloxidase cascade (Cerenius et al., 2008; Soderhall & Cerenius, 1998); total phenoloxidase activity was measured using an established protocol (Changsut et al., 2022; Fuess et al., 2016) which begins with a trypsin incubation step to convert any inactive prophenoloxidase into phenoloxidase. The resulting data therefore reflects the combined activity of inactive prophenoloxidase and active phenoloxidase in a sample, herein termed total phenoloxidase (TPO). Melanin concentration was also measured directly using established protocols (Changsut et al., 2022; Fuess et al., 2016). As melanin is a pigment rather than a protein, its concentration is standardized to dry tissue weight. Finally, we assessed the activity of antibacterial compounds in each sample (for this assay samples were all diluted to the lowest protein concentration) against *V. coralliilyticus*, capturing bacterial growth rate inhibition by coral extracts (Changsut et al., 2022). Corals produce a wide diversity of antibacterial compounds which may directly harm potential pathogens (Emery et al., 2024).

### 2.8 Statistical analyses: Experimental immune data

All statistical analyses for immune assays were conducted in R version 4.4.1 (Team, 2024). We began by assessing the impacts of environmental manipulation on symbiont density to confirm observed patterns of symbiont loss. Symbiont density was square root transformed to meet assumptions of normality, and then assessed in relation to temperature, nutrients, and their interaction using a linear mixed model (LMM). Genotype was fitted as a random effect to account for colony replication across treatments using the R package lme4 (Bates et al., 2015)

Immune data was analyzed to specifically assess the impacts of temperature and nutrient treatment (and the interaction thereof) on constitutive and induced immune responses. Prior to all analyses the R package mice (Buuren & Groothuis-Oudshoorn, 2011) was used to impute one missing catalase value using the pmm method with 5 imputed datasets and a maximum of 50 iterations. Initially, we conducted multivariate analyses of the impacts of our three factors (temperature, nutrients, immune challenge) on all immunological metrics (catalase, peroxidase, total phenoloxidase, melanin and antibacterial activity) and symbiont density data combined. Data was normalized and a Euclidean distance similarity matrix calculated. A PERMANOVA analysis was run using the R package vegan with the adonis2 function using the model: physiological metrics ∼ temperature * nutrients * immune Challenge (Martinez Arbizu, 2020; Oksanen et al., 2025). Genotype was not accounted for due to constraints of the adonis2 function. Post-hoc pairwise comparisons were conducted for significant terms using a custom function with the same parameters as the main model and corrected with a Bonferroni correction. To visualize significant effects, a principal component analysis of the data was run using the base R function prcomp and visualized using the R package factorextra with the fviz_pca_biplot function (Kassambara & Mundt, 2020). Principal component loadings and scores for the first through fourth components were then extracted and statistically analyzed for association with factors of interest. Associations were tested with a LMM with the formula: PC score ∼ temperature * nutrients * immune + genotype, where genotype is a random effect accounting for replication across treatments. Post-hoc analyses were conducted using the R package emmeans (Lenth, 2024) with Tukey p-value adjustments. Representative plots were constructed using ggplot2 (Wickham, 2016).

Following multivariate analyses, we examined each immunological metric independently using an LMM with the formula: immune metric ∼ temperature * nutrients * immune + genotype, where genotype is a random effect accounting for replication across treatments. Data was checked against appropriate statistical assumptions (normality, no outliers, etc.) prior to analyses and transformed when necessary. Post-hoc analyses were conducted using the R package emmeans with Tukey p-value adjustments. Representative plots were constructed using ggplot2 (Wickham, 2016).

## 3. Results

### 3.1 Modeling results: Disease susceptibility

SCTLD prevalence was most strongly influenced by the interactions between maximum temperature and maximum chlorophyll-a concentration the month prior to the survey, as well as between maximum chlorophyll-a concentration and three-month mean PAR prior to the survey. Disease probability increased with maximum temperature until maximum chlorophyll concentration exceeded ∼2 mg m^-3^, with colonies 50-60% more likely to develop SCTLD with each unit increase in temperature (Figure 3a). Around the mean regional maximum chlorophyll-a concentration (3.8 mg m^-3^), there was a slight negative relationship between maximum temperature and SCTLD prevalence, but this was not significantly different to zero until chlorophyll-a concentration exceeded ∼6 mg m^-3^. At that point, the probability of a colony having SCTLD declined substantially with maximum temperature. Disease probability declined with maximum chlorophyll-a concentration when the three-month mean PAR was below or equal to the FCR mean (52 einsteins m^-2^ s^-1^) but increased with chlorophyll-a concentration if three-month mean PAR was greater than the FCR mean (Figure 3b).

**Figure 3.**
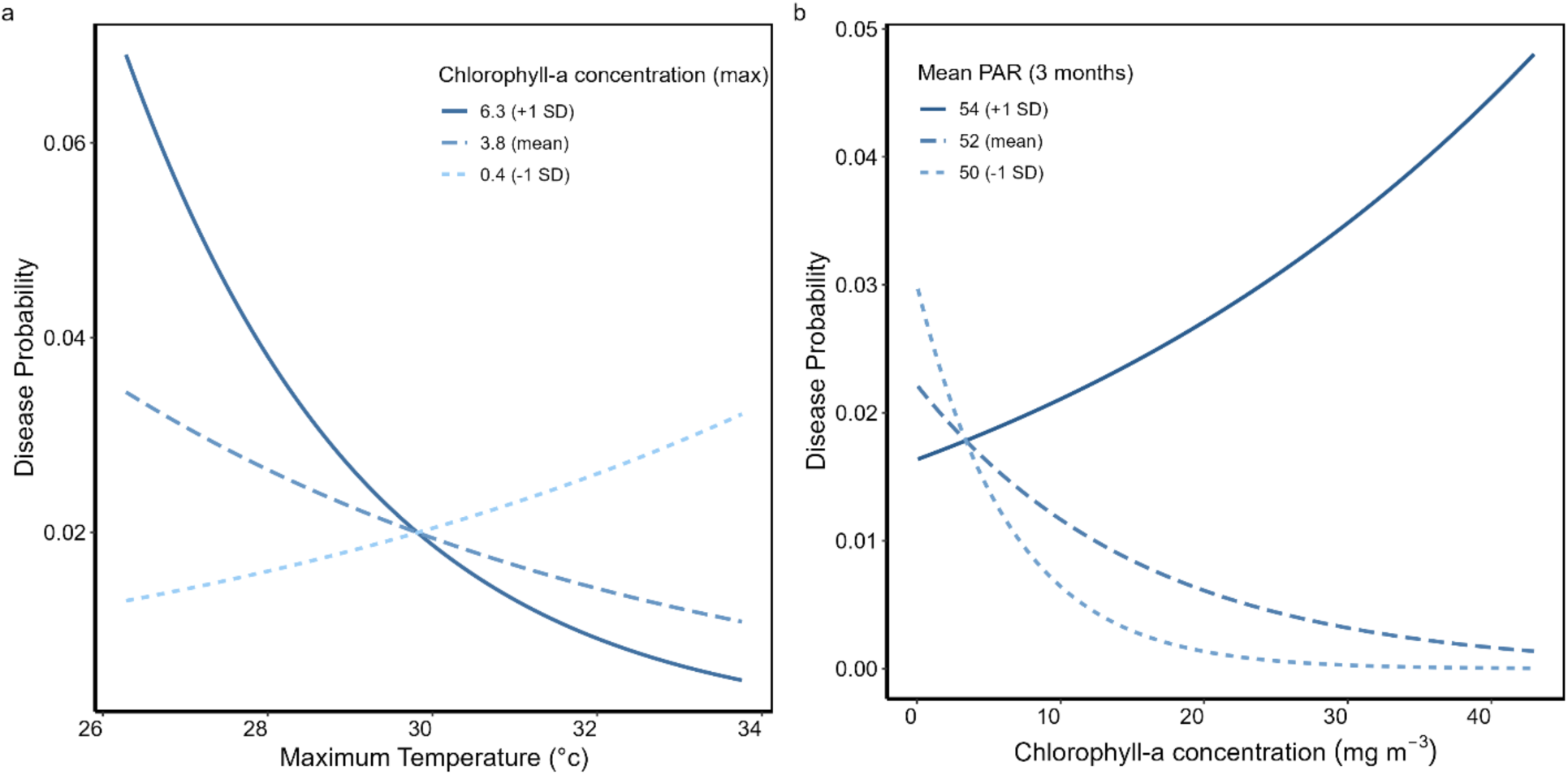
The greatest amount of variation in SCTLD prevalence was explained by the interaction between a) maximum temperature and maximum chlorophyll-a concentration, then b) maximum chlorophyll-a concentration and three-month mean PAR. The dotted line shows the concentration 1 standard below the mean, the dashed line the mean and the solid line 1 standard deviation above the mean value for a) chlorophyll-a concentration and b) mean PAR.

The probability of a colony having SCTLD also significantly increased with maximum PAR and declined with the mean temperature three months before surveying (GLMM; conditional R^2^ = 0.7, Marginal R^2^ = 0.1; Table 2). Colonies were 50% more likely to have SCTLD with every unit increase in maximum PAR and two times less likely to have SCTLD with every unit increase in three-month mean temperature. As indicated by the high conditional R^2^, survey location/time had a very large effect on SCTLD prevalence.

**Table 2.** Significant predictors of SCTLD prevalence (Note: maximum chlorophyll-a concentration retained due to presence in interactions) from GLMM. Estimate gives effect size on the logit scale. Negative estimate indicates a negative relationship between predictor and SCTLD prevalence. Bold font indicates significant p-values. *p < 0.05, **p < 0.001.

| Environmental predictor | Estimate | Std. Error | z value | p-value |
| --- | --- | --- | --- | --- |
| (Intercept) | -0.57 | 4.13 | -0.14 | 0.9 |
| Max temperature | 0.44 | 0.18 | 2.40 | **0.02\*** |
| Max PAR | 0.05 | 0.02 | 2.64 | **0.008\*\*** |
| 3-month mean temperature | -0.29 | 0.14 | -2.03 | **0.04\*** |
| Max Chl-a | 0.49 | 0.54 | 0.91 | 0.4 |
| 3-month mean PAR | -0.22 | 0.05 | -4.48 | **<0.001\*** |
| Max temp x Max Chl-a | -0.09 | 0.02 | -3.95 | **<0.001\*** |
| Max Chl-a x 3 month mean PAR | 0.04 | 0.01 | 4.58 | **<0.001\*** |

### 3.2 Modeling results: Disease severity

The probability of an *M. cavernosa* colony dying during the SCTLD outbreak significantly declined with maximum temperature (binomial GLM; R^2^ = 0.56; logit estimate = -0.82 ± 0.1 SE, p = 3.9 x 10^-11^), such that for every unit increase in maximum temperature, which averaged 31.4 °C (± 0.7 SD), colony mortality was three times less likely (Figure 4). Below 31.08 °C, the fitted GLM predicted there was a greater than 50% chance that an *M. cavernosa* colony would die. No other environmental predictor significantly affected mortality probability.

**Figure 4.**
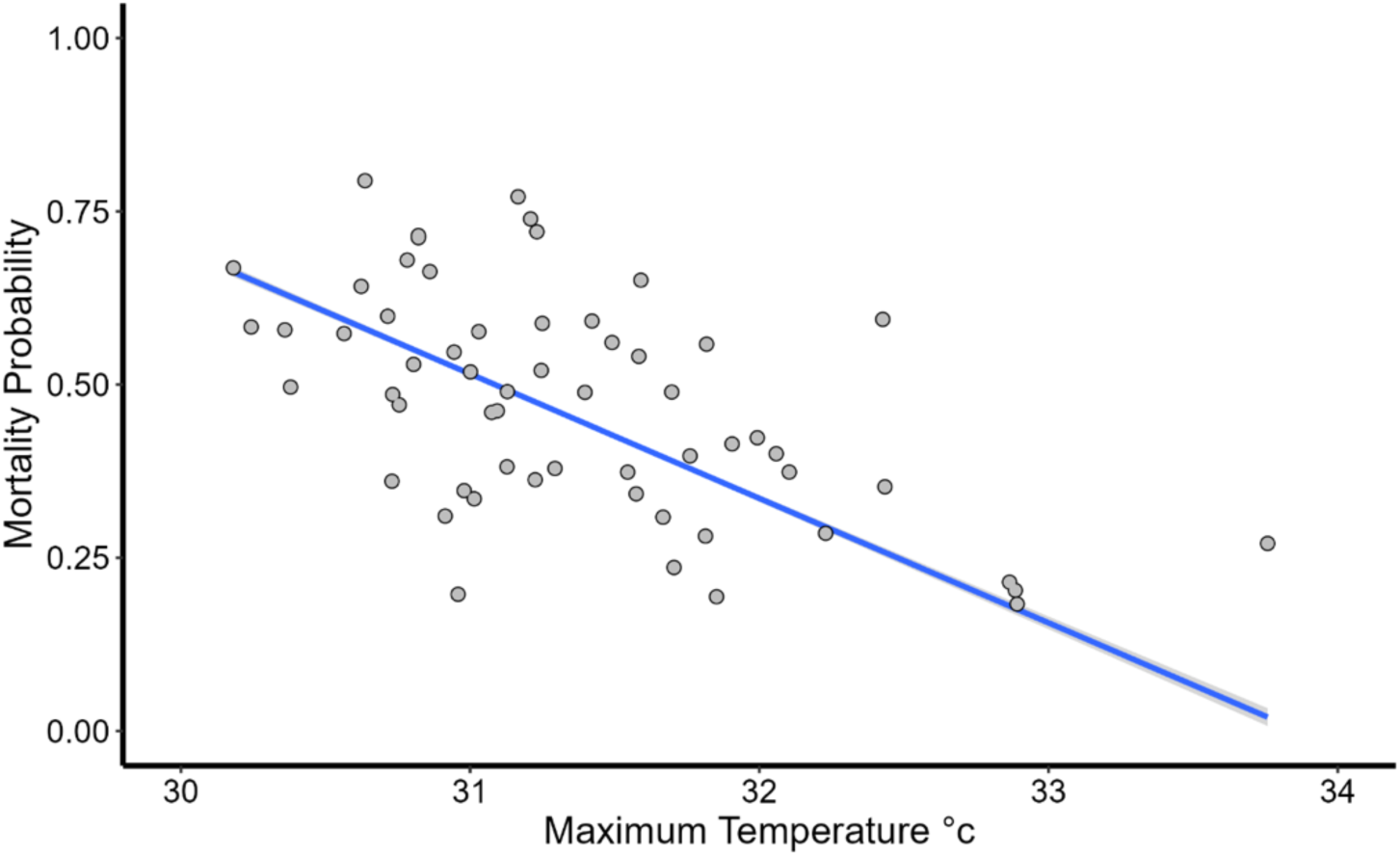
Modelled relationship between mortality probability for an *M. cavernosa* colony during the SCTLD outbreak (disease severity) and the maximum temperature in the two years between surveys (R^2^ = 0.56; estimate = -0.82 ± 0.1, p = 3.9 x 10^-11^). Blue line = mean mortality probability; grey points = model estimates for individual survey sites.

### 3.3 Experimental results: Symbiont density

Symbiont density varied significantly between temperature treatments (LMM; F = 40.99, p < 0.001) and marginally due to the interaction of temperature treatment and nutrients (LMM; F = 2.87, p = 0.06; Table S3). Symbiont density was generally higher in control colonies than heat-treated colonies (**Figure 5**). Furthermore, while not statistically significant, there was a notable trend wherein the difference between control and heat-treated colony symbiont density increased with nutrient levels.

**Figure 5.**
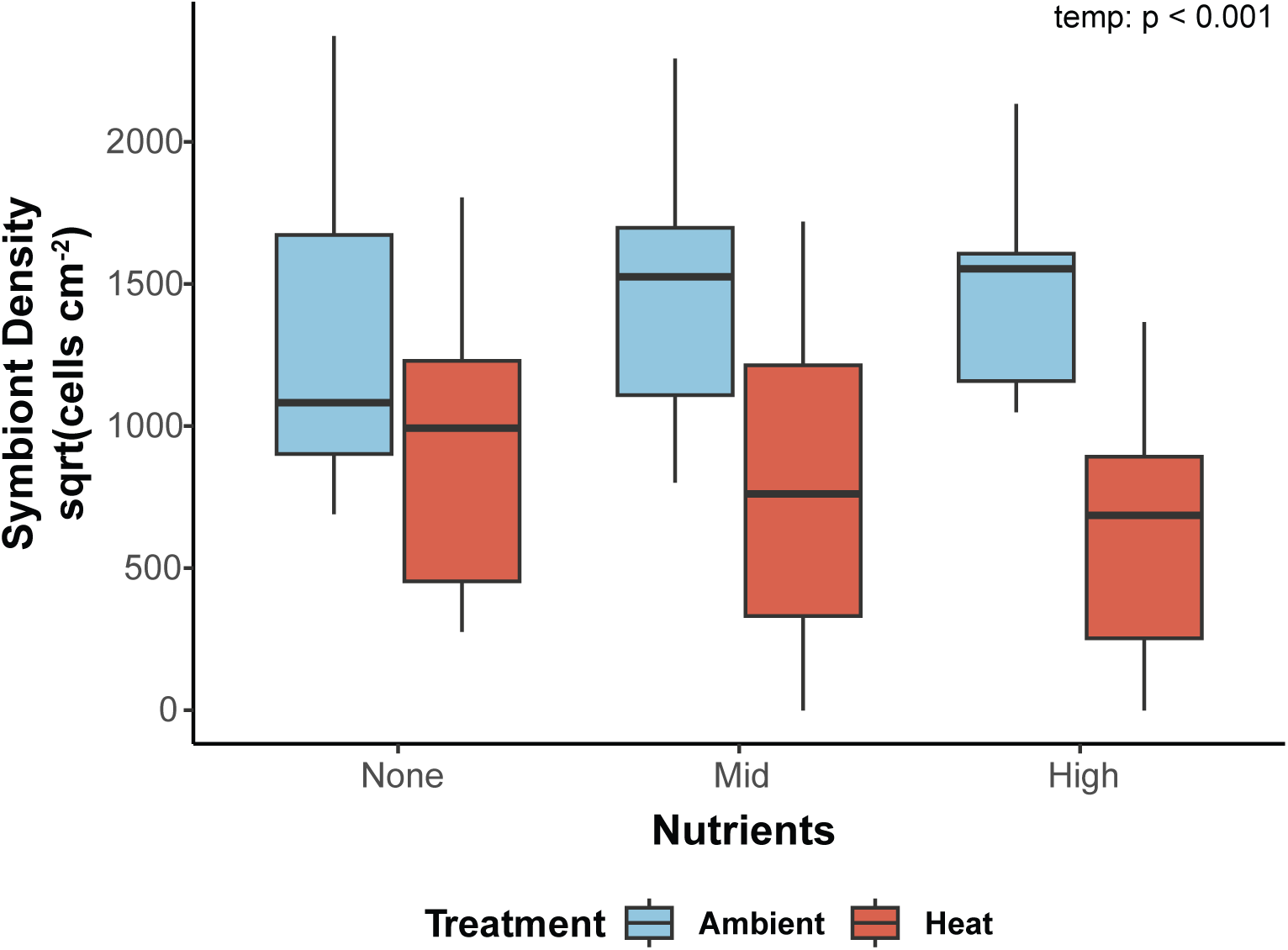
Box plot displaying square root transformed symbiont density data for colonies split by temperature treatment and nutrient level. Symbiont density was significantly higher in corals maintained at ambient temperatures (LMM; F = 40.99, p < 0.001) and marginally due to the interaction of temperature treatment and nutrients (LMM; F = 2.87, p = 0.06). Boxes are colored based on temperature treatment (blue = ambient: 28 °C, red = heat 32 °C). Mid nutrients = 0.01 mg/l ammonium, High nutrients = 0.05 mg/l ammonium. P values for significant model terms are listed.

### 3.4 Experimental results: Multivariate analyses

PERMANOVA analysis revealed strong impacts of temperature, nutrients, and their interaction on combined immune/symbiont metric data, but no impact of immune challenge (Table S4). Only 25% of the variance was explained by a factor or interaction of interest (PERMANOVA; residual R^2^ = 0.75). Temperature accounted for over 10% of the variance in immune metrics (PERMANOVA; R^2^ = 0.11, p = 0.001), whereas nutrients accounted for only about 4% of the variance (PERMANOVA; R^2^ = 0.04, p = 0.03). Finally, the combined impact of nutrients and temperature accounted for an additional 4.6% of the variance (PERMANOVA; R^2^ = 0. 0.046, p = 0.02). Pairwise analyses indicated significant differences between the high temp + mid nutrient groups (p = 0.02). The ambient temperature + mid nutrients treatment was significantly different from all heat treatment groups regardless of nutrient treatment (Table S5). Furthermore, the heat and mid nutrients treatment was significantly different from all ambient temperature treatments as well as the heat + high nutrient treatment (Table S5). Finally, the heat + no nutrient and heat + high nutrient treatments were significantly different from ambient temperature + mid nutrient and ambient + high treatments (Table S5).

Principal component analysis demonstrated clear clustering by temperature and nutrients, and some differentiation due to their combination (Figure 6). Principal components 1 through 4 explained a combined 80.33% of the variance (Table S6) and were tightly correlated with various immune and symbiont metrics (Table S6). PC1 explained roughly 28% of the variance and was strongly positively associated with symbiont density and all immune metrics except for catalase, which was negatively associated with PC1. PC1 was significantly different across temperature treatments (LMM: F = 56.4, p < 0.001, Table S7); specifically higher in ambient than heat treated fragments (Figure 7a). PC2 explained roughly 21% of the variance and was strongly positively associated with symbiont density, catalase and antibacterial activity, and melanin concentration. PC2 was significantly affected by the interaction of temperature and nutrients (LMM: F = 6.96, p = 0.002, Tables S7-8), with significant differences in immune metrics between ambient and heat treatments in the high nutrient group only (Figure 7b). PC3 explained an additional 18% of the variance and was strongly positively associated with catalase and total phenoloxidase but negatively associated with melanin concentration. PC3 varied significantly across nutrient concentrations (LMM; F = 9.28, p < 0.001, Tables S7-8), being negatively related with high nutrient treated corals compared to mid (Figure 7c). PC4 explained 13% of the variance and was positively correlated with antibacterial activity, total phenoloxidase and melanin concentration, and negatively associated with peroxidase activity and symbiont density. PC4 was significantly different due to the interaction between temperature and nutrients (LMM; F = 5.852, p = 0.005; Tables S7-S8), as well as nutrients and immune challenge (LMM; F = 4.729, p = 0.012; Tables S7-S8). Specifically, PC4 was significantly higher in colonies at high rather than ambient temperature under mid nutrients only (Figure 7d). Additionally, immune challenged fragments were significantly different in PC4 scores between mid and high nutrient treated individuals, with high nutrient immune challenged individuals having the greater PC4 scores than those that were immune challenged under mid conditions (Figure 7e). PC5 and PC6 explained 12% and 8% of the variance respectively (Table S6); neither was significantly associated with any factor of interest.

**Figure 6.**
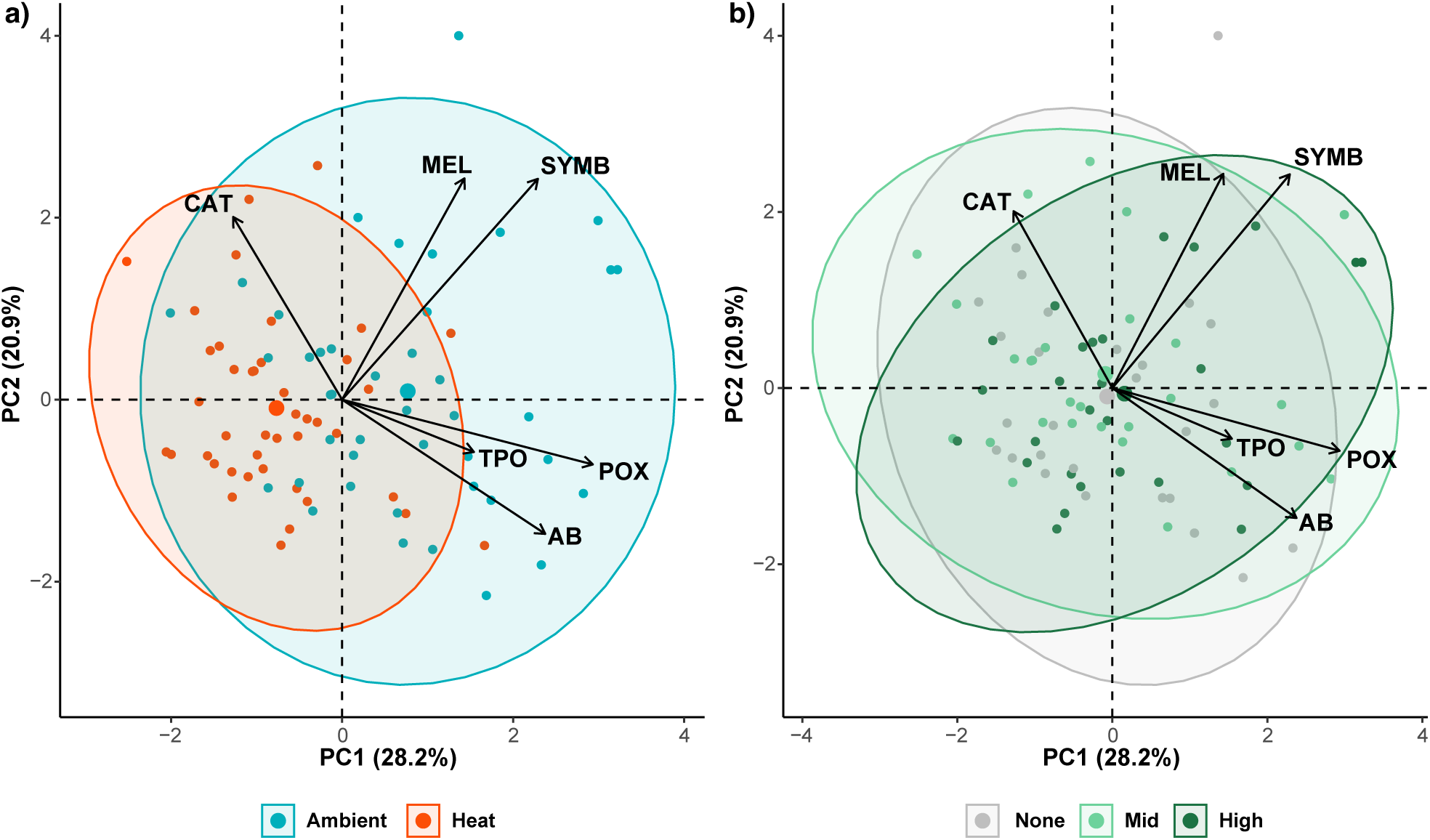
Biplot representation of the first two axes PCA analysis of combined symbiont and immune metric data. Plots are split based on groupings as follows: a) temperature treatment and b) nutrient treatment. In each case, points represent individual samples and are colored according to treatment group. Arrows depict eigenvector loadings of individual immune metrics, indicating direction in PC space and magnitude (i.e. length of line). Ellipses are drawn based on treatment groupings with 95% confidence intervals. PCA was visualized using the R package factorextra with the fviz_pca_biplot function.

**Figure 7.**
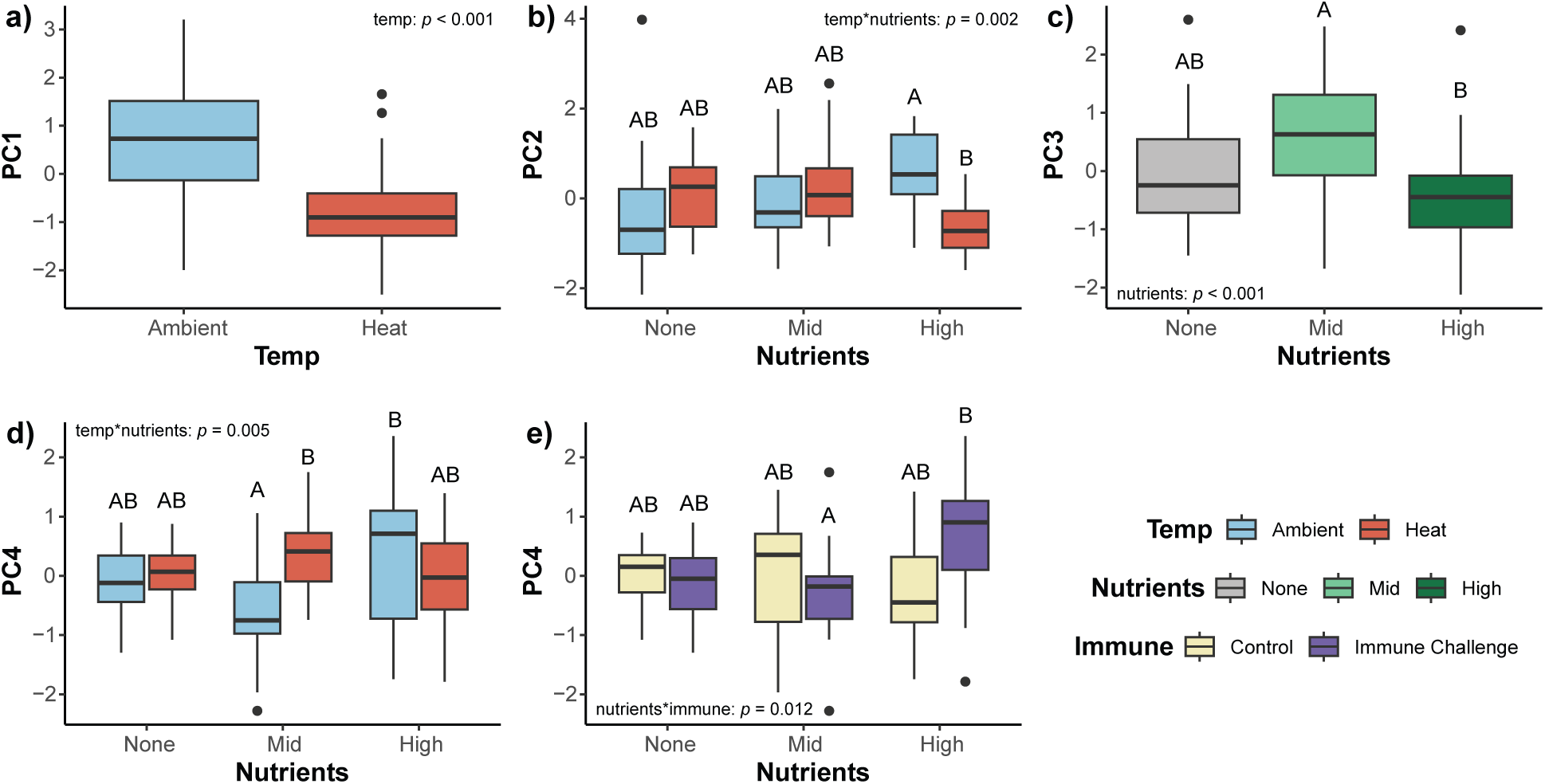
Boxplot representing significant variation in PC axis scores across temperature, nutrients, and immune challenge based on LMM results. Boxes are colored based on groupings according to legend. Letters represent statistically significant groups as detected by Tukey post-hoc tests (Full results shown in Table S8). P-values for significant model terms are listed. a) PC1, b) PC2, c) PC3, d-e) PC4.

### 3.5 Univariate Analyses of Immune Metrics

Univariate analyses revealed strong singular and interactive effects of temperature, nutrients, and immune challenge on baseline and induced immune activity (Table 3; Table S9; Figure 8). Increased temperature resulted in decreased baseline activity of peroxidase, total phenoloxidase, and antibacterial activity (Table 3; Figure 8b, c, e). Furthermore, the interaction of temperature and nutrient treatment significantly impacted melanin concentration (Table 3; Figure 8d); melanin concentration was significantly greater under high nutrient conditions compared to none and mid, but only at ambient temperature. Nutrient treatment also significantly impacted catalase activity generally and following immune challenge (Table 3; Figure 8a). Catalase activity was generally higher in mid nutrient conditions than high nutrient conditions (Tukey post-hoc, p = 0.007; Table S9) and was significantly higher in immune challenged corals under mid nutrient conditions compared to high nutrient conditions (Tukey post-hoc, p = 0.002; Table S9). Finally, immune challenge induced significant increases in antibacterial activity regardless of environmental conditions (LMM, p = 0.04; Table 3; Figure 8f).

**Figure 8.**
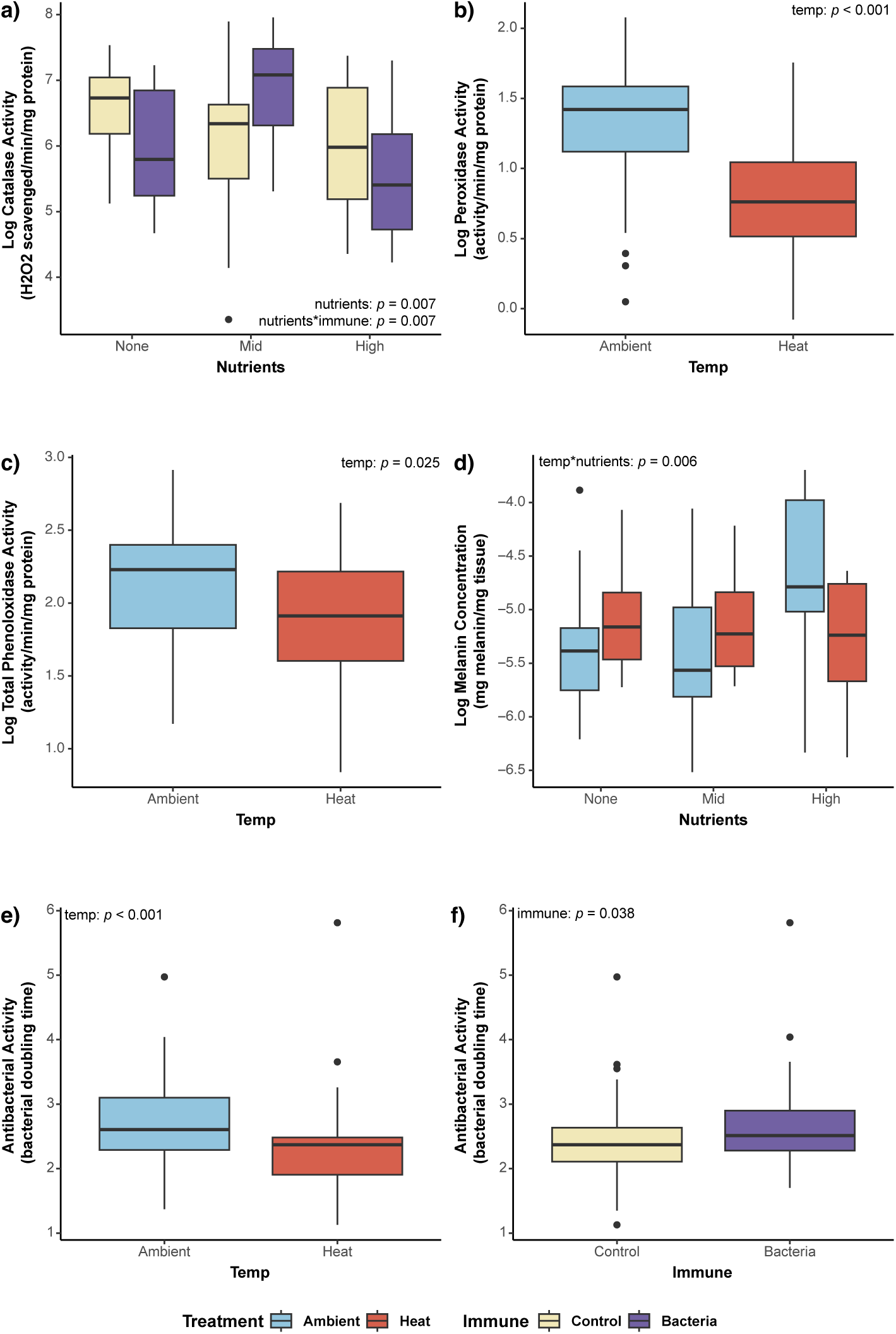
Boxplot representing significant variation in immune metrics across temperature, nutrients, and immune challenge based on LMM results. Normalized values are shown where appropriate. Boxes are colored based on groupings according to legend. Letters represent statistically significant groups as detected by Tukey post-hoc tests (Full results shown in Table S9). P-values for significant model terms are listed. a) log-normalized catalase activity, b) lognormalized peroxidase activity, c) log-normalized total phenoloxidase activity, d) log normalized melanin concentration, e-f) antibacterial activity

**Table 3.** LMM results of univariate analyses of variation in each immune metric due to temperature, nutrients, immune challenge, and their interactions. DF = degrees of freedom; Num = Numerator, Den = denominator. Bold font indicates significant p-values. *p < 0.05, **p < 0.001.Post-hoc test results for significant terms are found in Table S9.

| **Catalase** |  |  |  |  |  |  |
| --- | --- | --- | --- | --- | --- | --- |
| **Effect** | **Sum of Squares** | **Mean Squares** | **NumDF** | **DenDF** | **F value** | **p-value** |
| Temperature | 0.51 | 0.51 | 1 | 66 | 0.724 | 0.40 |
| Nutrients | 7.60 | 3.80 | 2 | 66 | 5.369 | **0.007\*** |
| Infection | 0.10 | 0.10 | 1 | 66 | 0.135 | 0.71 |
| Temperature: Nutrients | 0.28 | 0.14 | 2 | 66 | 0.197 | 0.82 |
| Temperature: Infection | 0.90 | 0.90 | 1 | 66 | 1.270 | 0.26 |
| Nutrients: Infection | 7.70 | 3.85 | 2 | 66 | 5.438 | **0.007\*** |
| Temperature: Nutrients: Infection | 2.66 | 1.33 | 2 | 66 | 1.880 | 0.16 |
| **Peroxidase** |  |  |  |  |  |  |
| **Effect** | **Sum of Squares** | **Mean Squares** | **NumDF** | **DenDF** | **F value** | **p-value** |
| Temperature | 5.22 | 5.22 | 1 | 66 | 33.71 | **<0.001\*\*** |
| Nutrients | 0.17 | 0.09 | 2 | 66 | 0.55 | 0.58 |
| Infection | 0.23 | 0.23 | 1 | 66 | 1.47 | 0.23 |
| Temperature: Nutrients | 0.86 | 0.43 | 2 | 66 | 2.76 | 0.07 |
| Temperature: Infection | 0.008 | 0.008 | 1 | 66 | 0.05 | 0.82 |
| Nutrients: Infection | 0.19 | 0.10 | 2 | 66 | 0.61 | 0.55 |
| Temperature: Nutrients: Infection | 0.17 | 0.08 | 2 | 66 | 0.54 | 0.59 |
| **Total Phenoloxidase** |  |  |  |  |  |  |
| **Effect** | **Sum of Squares** | **Mean Squares** | **NumDF** | **DenDF** | **F value** | **p-value** |
| Temperature | 0.95 | 0.95 | 1 | 66 | 5.25 | **0.03\*** |
| Nutrients | 0.78 | 0.39 | 2 | 66 | 2.16 | 0.12 |
| Infection | 0.09 | 0.09 | 1 | 66 | 0.48 | 0.49 |
| Temperature: Nutrients | 0.35 | 0.17 | 2 | 66 | 0.96 | 0.39 |
| Temperature: Infection | 0.01 | 0.01 | 1 | 66 | 0.07 | 0.78 |
| Nutrients: Infection | 0.36 | 0.18 | 2 | 66 | 0.98 | 0.38 |
| Temperature: Nutrients: Infection | 0.05 | 0.02 | 2 | 66 | 0.13 | 0.88 |
| **Melanin** |  |  |  |  |  |  |
| **Effect** | **Sum of Squares** | **Mean Squares** | **NumDF** | **DenDF** | **F value** | **p-value** |
| Temperature | 0.05 | 0.05 | 1 | 66 | 0.14 | 0.71 |
| Nutrients | 1.57 | 0.79 | 2 | 66 | 2.38 | 0.10 |
| Infection | 0.007 | 0.007 | 1 | 66 | 0.02 | 0.88 |
| Temperature: Nutrients | 3.69 | 1.84 | 2 | 66 | 5.58 | **0.006\*** |
| Temperature: Infection | 0.96 | 0.96 | 1 | 66 | 2.89 | 0.09 |
| Nutrients: Infection | 1.98 | 0.99 | 2 | 66 | 3.00 | 0.06 |
| Temperature: Nutrients: Infection | 0.27 | 0.14 | 2 | 66 | 0.41 | 0.66 |
| **Antibacterial activity** |  |  |  |  |  |  |
| **Effect** | **Sum of Squares** | **Mean Squares** | **NumDF** | **DenDF** | **F value** | **Pr(>F)** |
| Temperature | 2.79 | 2.79 | 1 | 63.98 | 12.36 | **<0.001\*\*** |
| Nutrients | 0.66 | 0.33 | 2 | 64.02 | 1.47 | 0.23 |
| Infection | 1.02 | 1.02 | 1 | 63.98 | 4.49 | **0.04\*** |
| Temperature: Nutrients | 0.64 | 0.27 | 2 | 64.09 | 1.20 | 0.31 |
| Temperature: Infection | 0.04 | 0.04 | 1 | 64.13 | 0.16 | 0.69 |
| Nutrients: Infection | 0.19 | 0.09 | 2 | 64.09 | 0.41 | 0.66 |
| Temperature: Nutrients: Infection | 0.17 | 0.08 | 2 | 64.02 | 0.37 | 0.69 |

## 4. Discussion

Our integrative combination of statistical modeling and experimental validation demonstrate clear impacts of environmental conditions on coral disease patterns and immunity. Modeling results demonstrate that variable environmental conditions, particularly temperature, nutrient enrichment, and light levels, significantly contribute to SCTLD prevalence and mortality in *Montastraea cavernosa*, which has also been found in previous coral disease outbreaks (Muller et al., 2008; Randall & van Woesik, 2015; Voss & Richardson, 2006). Furthermore, results from experimental validation studies indicate several of these factors, specifically temperature and nutrient enrichment, have impacts on host immunity. Furthermore, disparity between modeling and experimental results was observed at times, highlighting the complex associations between environmental factors, SCTLD dynamics, and host immunity which warrant further investigation.

### Temperature has complex impacts on SCTLD susceptibility, mortality, and host immunity

Elevated temperatures were associated with decreased coral mortality and SCTLD prevalence, as has been suggested elsewhere for SCTLD specifically (Palacio-Castro et al., 2025; Williams et al., 2021). This negative association between temperature and disease prevalence/mortality is starkly different than trends observed in previous coral disease outbreaks (van Woesik & Randall, 2017) wherein the negative effects of higher temperatures on host physiology are believed to facilitate synergistic interactions between temperature and disease (Lesser et al., 2007). This unique negative association between temperature and SCTLD has been hypothesized to be related to two factors. First, several studies have suggested that SCTLD initially affects Symbiodiniaceae before causing host tissue necrosis (e.g., Landsberg et al. 2020; Klein et al. 2024). Thus, high temperatures which induce bleaching or symbiont loss (as was demonstrated in this experiment) may mitigate SCTLD. Second, it is possible that high temperatures have a negative effect on the pathogen itself. Under ex-situ conditions, Palacio-Castro et al. (2025) found SCTLD contraction in another massive coral, *Orbicella faveolata,* declined at high temperature despite no bleaching, suggesting temperature directly suppressed pathogenicity. Indeed, particularly when combined with our experimental results, our modeling data strongly support an important role of secondary environmental effects on either the SCTLD pathogen or Symbiodiniaceae in determining synergistic disease outcomes. Further study investigating the mechanisms of these secondary effects is essential to accurately modeling SCTLD outcomes.

In contrast to patterns observed in modeling, our experimental results provide compelling evidence that prolonged exposure to elevated temperatures has significant negative effects on host immunity, as has been widely hypothesized and observed (Lesser et al., 2007). Both multivariate and univariate modeling revealed strong effects of elevated temperatures on all three measured metrics of coral immunity (antioxidant activity, melanin synthesis, and antibacterial activity). Antioxidant activity patterns following heat stress were mixed; catalase activity was increased whereas peroxidase activity was decreased. These patterns are reflective of the dual role of antioxidant activity in protecting the host against ROS produced during both stress and immune response (Emery et al., 2024). Although ROS are essential physiological regulators under normal conditions, at high concentrations they can induce oxidative stress that can cause significant damage to DNA and proteins (Hayat, 2015). Specifically, in corals, temperature and light stress can stimulate the production of extremely high concentrations of ROS (Hansel & Diaz, 2021). The inverse relationship between two antioxidants measured in this experiment can also likely be explained by diversification of function amongst coral antioxidants. Catalase has been observed as the most responsive coral antioxidant to elevated temperatures in other stony corals, in agreement with our data (Krueger et al., 2015). In contrast, coral peroxidases have been strongly documented for their potential roles in immunity. In the octocoral, *Gorgonia ventalina*, peroxidases are inducible following pathogen stress and have direct antifungal activities (Mydlarz & Harvell, 2007). Thus, while the upregulation of catalase observed here is likely indicative of coral responses to, and perhaps coping with, higher temperatures, the downregulation of peroxidase is more likely reflective of suppressed immune responses.

Both components of the melanin synthesis cascade (total phenoloxidase and melanin concentration) were significantly suppressed in corals exposed to prolonged elevated temperature. Previous studies have highlighted the importance for both total phenoloxidase activity and melanin concentration for direct defense response (Palmer et al., 2008) and have shown associations between levels of these responses and coral disease susceptibility (Palmer et al., 2010; Palmer, Bythell, et al., 2011; Pinzon et al., 2014). Furthermore, past evidence has highlighted the negative impacts of elevated temperatures and/or bleaching on components of the melanin synthesis cascade, though these trends may be species specific (Palmer, Bythell, et al., 2011; van de Water et al., 2016) Notably, our study is among the first to confirm temperature-mediated suppression of the melanin synthesis cascade in a laboratory setting. This adds clarity to previous observations from field-based studies (Palmer, Bythell, et al., 2011; van de Water et al., 2016) and provides support that prolonged elevated temperatures have strong negative effects on coral immunity and potentially disease susceptibility.

Finally, we also observed suppression of antibacterial activity in response to heat treatment, which likely further reduces coral immunity and exacerbates general disease susceptibility in *M. cavernosa*. Our laboratory results also highlight the importance of antibacterial compounds in defense against pathogens, as antibacterial activity increased, irrespective of heat or nutrient treatment, following pathogen exposure. Further, antibacterial activity is known to be important for SCTLD defense; the application of natural antibacterial producing probiotics or antibacterial compounds has been shown to be an effective treatment for SCTLD (Aeby et al., 2019; Forrester et al., 2022; Neely et al., 2020; Shilling et al., 2021; Ushijima et al., 2023; Walker et al., 2021). Thus, the suppression of antibacterial activity due to prolonged temperature could have significant impacts on coral disease susceptibility.

It should be noted that we observed contrasting effects of temperature on coral health across our modeling and laboratory settings. These differences suggest the impact of environmental conditions on the unknown pathogen driving SCTLD also strongly impact disease dynamics. Thus, it is essential to take an integrative approach incorporating host and pathogen when studying the environmental effects on diseases. It is clear that accurately predicting future outbreaks of SCTLD and similar diseases will be difficult without the identification of a pathogen and integration of its unique biology into modeling efforts.

### Nutrient enrichment has complex impacts on coral immunity

Modeling and ex-situ studies both found an effect of nutrients on disease dynamics and immunity, respectively. In modeling, the nutrient proxy, chlorophyll-a, positively influenced SCTLD susceptibility, presumably by either directly compromising *M. cavernosa* health or influencing the Symbiodiniaceae (Bell et al., 2014; Lapointe et al., 2019). Meanwhile, variations in ammonium concentration modified both baseline and induced immune responses in laboratory settings. Corals kept at moderate ammonium levels (0.01 mg/L) had higher levels of catalase activity and total phenoloxidase, than those kept in absence of ammonium. The induced immune response (measured via pathogen-injected corals) also appeared to be modulated by moderate ammonium levels – catalase activity was increased in these samples. In contrast, fragments reared under high ammonium concentrations had greater antibacterial activity and melanin synthesis activity (both total phenoloxidase and melanin). Previous studies have demonstrated indirect negative effects of ammonium on coral immunity, which are presumably modulated by changes in symbiont density (Fuess et al., 2020). Additionally, other studies have highlighted suppression of phenoloxidase activity following nutrient enrichment with fertilizers (Dougan et al., 2020). However, some studies have failed to find any impacts of nitrate enrichment on catalase activity, though they did document increased activity under multiple stressor conditions (Higuchi et al., 2015) The lack of consistent methods across limited studies makes it difficult to draw any sweeping conclusions regarding the impacts of nutrient enrichment on coral immunity. However, our results do highlight significant nuance in nutrient-immune relationship, suggesting there is an optimal balance of nutrients which promotes coral immunity, as has been shown in other studies (Becker et al. 2021; Shantz et al 2023).

### Both SCTLD susceptibility and host coral immunity are significantly impacted by the interaction of temperature and nutrient enrichment

In addition to main effects of temperature and nutrients, we also observed strong interactive effects of these two environmental conditions on both SCTLD susceptibility in the field and on constitutive coral immunity in the laboratory. SCTLD prevalence in *M. cavernosa* was most strongly influenced by the interactive effects of temperature and chlorophyll-a, and of photosynthetically active radiation (PAR) and chlorophyll-a. Under low chlorophyll-a concentrations, an *M. cavernosa* colony was 50-60% more likely to have SCTLD with every 1° C increase in maximum temperature, a relationship frequently observed with other coral diseases (Muller et al., 2018; Randall & van Woesik, 2015; Sato et al., 2009). However, when chlorophyll-a was high, this relationship flipped, and SCTLD prevalence was highest at lower temperature, particularly below 28 °C. These results do counter Muller et al. (2020), who found no effect of temperature or chlorophyll on the initial SCTLD outbreak, likely due to earlier temporal scale and the confounding effect of coral community composition rather than the single species approach taken here. Chlorophyll-a has increased significantly since the 1980s in Florida and has been linked to previous disease outbreaks (Lapointe et al., 2019). High levels of chlorophyll-a are predominant across Florida (Lapointe et al., 2019) and, combined with our results, are likely to explain the negative associations between temperature and SCTLD observed in other studies(Williams et al., 2021)

We also observed complex interactive effects of elevated temperature and ammonium on constitutive coral immunity in our laboratory experiments. The combination of high ammonium and high temperature significantly reduced catalase and antibacterial activity. In contrast, corals exposed to elevated levels of one stressor (either high temperature or high ammonium with the other factor at ambient/control levels) had higher antibacterial and melanin synthesis activity than those exposed to moderate nutrients and ambient temperatures. Additionally, melanin was highest under high ammonium at ambient temperature only. This suggests that while exposure to a single environmental stressor may elicit a stress or even priming response, combinations of stressors result in general immune suppression. Previous studies have demonstrated that single stressors may elicit an increased coral immune response (Eliachar et al., 2022; Sharoni et al., 2025; van de Water et al., 2018), though these may be short-lived stress responses. Prior documentation of responses to multiple stressors is limited, revealing mixed and potentially species specific patterns; some studies document enhanced immune responses in the presence of multiple stressors, whereas others document antagonistic interactions (Dilworth et al., 2024; van de Water et al., 2015). Overall, the interactive effects observed in both modeling and laboratory experiments are complex and warrant further study.

While only tested in our modeling study, the additive effect of high chlorophyll-a concentration and prolonged PAR intensity also significantly increased SCTLD probability. Light intensity is frequently reported as having a compounding effect with temperature on coral bleaching (Morris et al. 2019) and disease (Ban et al., 2014; Randall & van Woesik, 2015; Sato et al., 2009) by increasing oxidative stress within the coral colony. However, the interactive effect of chlorophyll or nutrients and light on disease is less studied. Elevated nitrogen concentrations have been shown to lower resistance to bleaching caused by light stress through phosphate starvation (Wiedenmann et al. 2013). Furthermore, algal symbiont density, which increases with chlorophyll concentration, has been correlated with oxidative stress (Cunning & Baker, 2012). It stands to reason that either mechanism, increased oxidative stress or phosphate starvation, likely enhanced disease susceptibility. For corals reefs to persist, improved understanding of how globally rising temperature and local environmental conditions interact to modulate coral immunity and future disease outbreaks is essential.

### Conclusions

Herein we demonstrate through combined statistical modeling of field observations and experimental validation that environmental conditions, particularly nutrient concentrations and temperature, have significant effects on SCTLD dynamics and host immunity. We experimentally validated that conditions largely considered “stressful” to corals impact immunity. It is important to note however that the impacts are disparate across stressors; temperature largely affects baseline immune responses whereas nutrients impact induced responses. Furthermore, our observation of the disparate effects of temperature in modeling and experimentation highlights the pressing need to identify and incorporate pathogen biology into our understanding of environment-disease associations. This point is epitomized by the contrast between the conventional view of coral disease dynamics and the identified effect of temperature on the mortality rate during the SCTLD outbreak in Florida. Future studies should delve further into the complexities described herein and incorporate knowledge gained into robust models to predict the future of coral disease dynamics.

## Acknowledgements

We would like to thank all CREMP, SECREMP and DRM researchers who collected field data, NASA and NOAA for satellite data and the Southeast Florida Reef Tract Water Quality Assessment Project and the Southeast Environmental Research Center for collecting in situ water quality data. Special thanks to the Coral Reef Restoration Assessment and Monitoring lab at Nova Southeastern University for supplying coral colonies. We would also like to thank the Florida Department of Environmental Protection Florida’s Coral Reef Resilience Program Grant award C40115 and NOAA’s Coral Reef Conservation Program award NA23NOS4690272 for funding this project.

## References

Aeby, G. S., Howells, E., Work, T., Abrego, D., Williams, G. J., Wedding, L. M., Caldwell, J. M., Moritsch, M., & Burt, J. A. (2020). Localized outbreaks of coral disease on Arabian reefs are linked to extreme temperatures and environmental stressors. Coral Reefs, 39(3), 829–846. 10.1007/s00338-020-01928-4

Aeby, G. S., Ushijima, B., Campbell, J. E., Jones, S., Williams, G. J., Meyer, J. L., Häse, C., & Paull, V. J. (2019). Pathogenesis of a Tissue Loss Disease Affecting Multiple Species of Corals Along the Florida Reef Tract. Frontiers in Marine Science, 6. 10.3389/fmars.2019.00678

Alvarez-Filip, L., Estrada-Saldivar, N., Perez-Cervantes, E., Molina-Hernandez, A., & Gonzalez-Barrios, F. J. (2019). A rapid spread of the stony coral tissue loss disease outbreak in the Mexican Caribbean. PeerJ, 7, e8069. 10.7717/peerj.8069

Alvarez-Filip, L., Gonzalez-Barrios, F. J., Perez-Cervantes, E., Molina-Hernandez, A., & Estrada-Saldivar, N. (2022). Stony coral tissue loss disease decimated Caribbean coral populations and reshaped reef functionality. Commun Biol, 5(1), 440. 10.1038/s42003-022-03398-6

Ban, S. S., Graham, N. A., & Connolly, S. R. (2014). Evidence for multiple stressor interactions and effects on coral reefs. Glob Chang Biol, 20(3), 681–697. 10.1111/gcb.12453

Barris, B. N., Shields, J. D., Small, H. J., Huchin-Mian, J. P., O’Leary, P., Shawver, J. V., Glenn, R. P., & Pugh, T. L. (2018). Laboratory studies on the effect of temperature on epizootic shell disease in the American lobster, Homarus americanus. Bulletin of Marine Science, 94(3), 887–902. 10.5343/bms.2017.1148

Bates, D., Mächler, M., Bolker, B., & Walker, S. (2015). Fitting Linear Mixed-Effects Models Usinglme4. Journal of Statistical Software, 67(1). 10.18637/jss.v067.i01

Belasen, A. M., Bletz, M. C., Leite, D. d. S., Toledo, L. F., & James, T. Y. (2019). Long-Term Habitat Fragmentation Is Associated With Reduced MHC IIB Diversity and Increased Infections in Amphibian Hosts. Frontiers in Ecology and Evolution, 6. 10.3389/fevo.2018.00236

Bell, P. R., Elmetri, I., & Lapointe, B. E. (2014). Evidence of large-scale chronic eutrophication in the Great Barrier Reef: quantification of chlorophyll a thresholds for sustaining coral reef communities. Ambio, 43(3), 361–376. 10.1007/s13280-013-0443-1

Berkhout, B. W., Budria, A., Thieltges, D. W., & Slabbekoorn, H. (2023). Anthropogenic noise pollution and wildlife diseases. Trends Parasitol, 39(3), 181–190. 10.1016/j.pt.2022.12.002

Brooks, M. E., Kristensen, K., Benthem, K. J. v., Magnusson, A., Berg, C. W., Nielsen, A., Skaug, H. J., Mächler, M., & Bolker, B. M. (2017). glmmTMB Balances Speed and Flexibility Among Packages for Zero-inflated Generalized Linear Mixed Modeling. The R Journal, 9(2). 10.32614/rj-2017-066

Bruno, J. F., Selig, E. R., Casey, K. S., Page, C. A., Willis, B. L., Harvell, C. D., Sweatman, H., & Melendy, A. M. (2007). Thermal stress and coral cover as drivers of coral disease outbreaks. PLoS Biol, 5(6), e124. 10.1371/journal.pbio.0050124

Burge, C. A., Mark Eakin, C., Friedman, C. S., Froelich, B., Hershberger, P. K., Hofmann, E. E., Petes, L. E., Prager, K. C., Weil, E., Willis, B. L., Ford, S. E., & Harvell, C. D. (2014). Climate change influences on marine infectious diseases: implications for management and society. Ann Rev Mar Sci, 6, 249–277. 10.1146/annurev-marine-010213-135029

Burke, S., Pottier, P., Lagisz, M., Macartney, E. L., Ainsworth, T., Drobniak, S. M., & Nakagawa, S. (2023). The impact of rising temperatures on the prevalence of coral diseases and its predictability: A global meta-analysis. Ecol Lett, 26(8), 1466–1481. 10.1111/ele.14266

Buuren, S. v., & Groothuis-Oudshoorn, K. (2011). mice: Multivariate Imputation by Chained Equations inR. Journal of Statistical Software, 45(3). 10.18637/jss.v045.i03

Caldwell, J. M., Liu, G., Geiger, E., Heron, S. F., Eakin, C. M., De La Cour, J., Greene, A., Raymundo, L., Dryden, J., Schlaff, A., Stella, J. S., Kindinger, T. L., Couch, C. S., Fenner, D., Hoot, W., Manzello, D., & Donahue, M. J. (2024). Multi-Factor Coral Disease Risk: A new product for early warning and management. Ecol Appl, 34(4), e2961. 10.1002/eap.2961

Carmignani, A., Skrzypek, G., Brooker, R. M., Meekan, M. G., Chase, T. J., Shantz, A. A., & Barneche, D. R. (2025). The relationship between nutrient supply from resident fishes and the growth, condition, and thermal tolerance of corals. Coral Reefs, 44(5), 1815–1837. 10.1007/s00338-025-02680-3

Cerenius, L., Lee, B. L., & Soderhall, K. (2008). The proPO-system: pros and cons for its role in invertebrate immunity. Trends Immunol, 29(6), 263–271. 10.1016/j.it.2008.02.009

Changsut, I., Womack, H. R., Shickle, A., Sharp, K. H., & Fuess, L. E. (2022). Variation in symbiont density is linked to changes in constitutive immunity in the facultatively symbiotic coral, Astrangia poculata. Biol Lett, 18(11), 20220273. 10.1098/rsbl.2022.0273

Cheng, T. L., Reichard, J. D., Coleman, J. T. H., Weller, T. J., Thogmartin, W. E., Reichert, B. E., Bennett, A. B., Broders, H. G., Campbell, J., Etchison, K., Feller, D. J., Geboy, R., Hemberger, T., Herzog, C., Hicks, A. C., Houghton, S., Humber, J., Kath, J. A., King, R. A., Frick, W. F. (2021). The scope and severity of white-nose syndrome on hibernating bats in North America. Conserv Biol, 35(5), 1586–1597. 10.1111/cobi.13739

Cohen, J. M., Sauer, E. L., Santiago, O., Spencer, S., & Rohr, J. R. (2020). Divergent impacts of warming weather on wildlife disease risk across climates. Science, 370(6519). 10.1126/science.abb1702

Cunning, R., & Baker, A. C. (2012). Excess algal symbionts increase the susceptibility of reef corals to bleaching. Nature Climate Change, 3(3), 259–262. 10.1038/nclimate1711

Dahlgren, C., Pizarro, V., Sherman, K., Greene, W., & Oliver, J. (2021). Spatial and Temporal Patterns of Stony Coral Tissue Loss Disease Outbreaks in The Bahamas. Frontiers in Marine Science, 8. 10.3389/fmars.2021.682114

Daszak, P., Cunningham, A. A., & Hyatt, A. D. (2000). Emerging infectious diseases of wildlife--threats to biodiversity and human health. Science, 287(5452), 443–449.

Dilworth, J., Million, W. C., Ruggeri, M., Hall, E. R., Dungan, A. M., Muller, E. M., & Kenkel, C. D. (2024). Synergistic response to climate stressors in coral is associated with genotypic variation in baseline expression. Proc Biol Sci, 291(2019), 20232447. 10.1098/rspb.2023.2447

Dortch, Q. (1990). The Interaction between Ammonium and Nitrate Uptake in Phytoplankton. Marine Ecology Progress Series, 61(1-2), 183–201. DOI 10.3354/meps061183

Dougan, K. E., Ladd, M. C., Fuchs, C., Vega Thurber, R., Burkepile, D. E., & Rodriguez-Lanetty, M. (2020). Nutrient Pollution and Predation Differentially Affect Innate Immune Pathways in the Coral Porites porites. Frontiers in Marine Science, 7. 10.3389/fmars.2020.563865

Eisenlord, M. E., Groner, M. L., Yoshioka, R. M., Elliott, J., Maynard, J., Fradkin, S., Turner, M., Pyne, K., Rivlin, N., van Hooidonk, R., & Harvell, C. D. (2016). Ochre star mortality during the 2014 wasting disease epizootic: role of population size structure and temperature. Philos Trans R Soc Lond B Biol Sci, 371(1689). 10.1098/rstb.2015.0212

Eliachar, S., Snyder, G. A., Barkan, S. K., Talice, S., Otolenghi, A., Jaimes-Becerra, A., Sharoni, T., Sultan, E., Hadad, U., Levy, O., Moran, Y., Gershoni-Yahalom, O., Traylor-Knowles, N., & Rosental, B. (2022). Heat stress increases immune cell function in Hexacorallia. Front Immunol, 13, 1016097. 10.3389/fimmu.2022.1016097

Emery, M., Gutierrez-Andrade, D., Changsut, I., Swain, H., Fuess, L., & Mydlarz, L. (2024). Immune System Components in Cnidarians. In. 10.1016/B978-0-128-24465-4.00122-8

Fisher, M. C., & Garner, T. W. J. (2020). Chytrid fungi and global amphibian declines. Nat Rev Microbiol, 18(6), 332–343. 10.1038/s41579-020-0335-x

Forrester, G. E., Arton, L., Horton, A., Nickles, K., & Forrester, L. M. (2022). Antibiotic Treatment Ameliorates the Impact of Stony Coral Tissue Loss Disease (SCTLD) on Coral Communities. Frontiers in Marine Science, 9. 10.3389/fmars.2022.859740

Fraser, R. N. (1998). Hyperspectral remote sensing of turbidity and chlorophyll a among Nebraska Sand Hills lakes. International journal of remote sensing, 19(8), 1579–1589.

Fuess, L. E., Palacio-Castro, A. M., Butler, C. C., Baker, A. C., & Mydlarz, L. D. (2020). Increased Algal Symbiont Density Reduces Host Immunity in a Threatened Caribbean Coral Species, Orbicella faveolata. Frontiers in Ecology and Evolution, 8. 10.3389/fevo.2020.572942

Fuess, L. E., Pinzon C, J. H., Weil, E., & Mydlarz, L. D. (2016). Associations between transcriptional changes and protein phenotypes provide insights into immune regulation in corals. Dev Comp Immunol, 62, 17–28. 10.1016/j.dci.2016.04.017

Gardner, S. G., Raina, J. B., Ralph, P. J., & Petrou, K. (2017). Reactive oxygen species (ROS) and dimethylated sulphur compounds in coral explants under acute thermal stress. J Exp Biol, 220(Pt 10), 1787–1791. 10.1242/jeb.153049

Hansel, C. M., & Diaz, J. M. (2021). Production of Extracellular Reactive Oxygen Species by Marine Biota. Ann Rev Mar Sci, 13, 177–200. 10.1146/annurev-marine-041320-102550

Hartig, F. (2017). DHARMa: residual diagnostics for hierarchical (multi-level/mixed) regression models R package. In

Harvell, C. D., Kim, K., Burkholder, J. M., Colwell, R. R., Epstein, P. R., Grimes, D. J., Hofmann, E. E., Lipp, E. K., Osterhaus, A. D., Overstreet, R. M., Porter, J. W., Smith, G. W., & Vasta, G. R. (1999). Emerging marine diseases--climate links and anthropogenic factors. Science, 285(5433), 1505–1510. 10.1126/science.285.5433.1505

Hayat, M. A. (2015). Introduction to Autophagy. In Autophagy: Cancer, Other Pathologies, Inflammation, Immunity, Infection, and Aging (pp. 1–53). 10.1016/b978-0-12-801043-3.00001-7

Hayes, N. K., Walton, C. J., & Gilliam, D. S. (2022). Tissue loss disease outbreak significantly alters the Southeast Florida stony coral assemblage. Frontiers in Marine Science, 9. 10.3389/fmars.2022.975894

Hewson, I., Button, J. B., Gudenkauf, B. M., Miner, B., Newton, A. L., Gaydos, J. K., Wynne, J., Groves, C. L., Hendler, G., Murray, M., Fradkin, S., Breitbart, M., Fahsbender, E., Lafferty, K. D., Kilpatrick, A. M., Miner, C. M., Raimondi, P., Lahner, L., Friedman, C. S., . . . Harvell, C. D. (2014). Densovirus associated with sea-star wasting disease and mass mortality. Proc Natl Acad Sci U S A, 111(48), 17278–17283. 10.1073/pnas.1416625111

Higuchi, T., Yuyama, I., & Nakamura, T. (2015). The combined effects of nitrate with high temperature and high light intensity on coral bleaching and antioxidant enzyme activities. Regional Studies in Marine Science, 2, 27–31. 10.1016/j.rsma.2015.08.012

Jones, N. P., Figueiredo, J., & Gilliam, D. S. (2020). Thermal stress-related spatiotemporal variations in high-latitude coral reef benthic communities. Coral Reefs, 39(6), 1661–1673. 10.1007/s00338-020-01994-8

Jones, N. P., & Gilliam, D. S. (2024). Temperature and local anthropogenic pressures limit stony coral assemblage viability in southeast Florida. Mar Pollut Bull, 200, 116098. 10.1016/j.marpolbul.2024.116098

Jones, N. P., Kabay, L., Semon Lunz, K., & Gilliam, D. S. (2021). Temperature stress and disease drives the extirpation of the threatened pillar coral, Dendrogyra cylindrus, in southeast Florida. Sci Rep, 11(1), 14113. 10.1038/s41598-021-93111-0

Kassambara, A., & Mundt, F. (2020). Factoextra: Extract and Visualize the Results of Multivariate Data Analyses. In https://CRAN.R-project.org/package=factoextra

Klinges, J. G., Patel, S. H., Duke, W. C., Muller, E. M., & Vega Thurber, R. L. (2022). Phosphate enrichment induces increased dominance of the parasite Aquarickettsia in the coral Acropora cervicornis. FEMS Microbiol Ecol, 98(2). 10.1093/femsec/fiac013

Krueger, T., Hawkins, T. D., Becker, S., Pontasch, S., Dove, S., Hoegh-Guldberg, O., Leggat, W., Fisher, P. L., & Davy, S. K. (2015). Differential coral bleaching-Contrasting the activity and response of enzymatic antioxidants in symbiotic partners under thermal stress. Comp Biochem Physiol A Mol Integr Physiol, 190, 15–25. 10.1016/j.cbpa.2015.08.012

Lapointe, B. E., Brewton, R. A., Herren, L. W., Porter, J. W., & Hu, C. (2019). Nitrogen enrichment, altered stoichiometry, and coral reef decline at Looe Key, Florida Keys, USA: a 3-decade study. Marine Biology, 166(8). 10.1007/s00227-019-3538-9

Lenth, R. (2024). emmeans: Estimated Marginal Means, aka Least-Squares Means. In (Version R package version 1.10.4)

Lesser, M. P., Bythell, J. C., Gates, R. D., Johnstone, R. W., & Hoegh-Guldberg, O. (2007). Are infectious diseases really killing corals? Alternative interpretations of the experimental and ecological data. Journal of Experimental Marine Biology and Ecology, 346(1-2), 36–44. 10.1016/j.jembe.2007.02.015

Liaw, A., & Wiener, M. (2002). Classification and regression by randomForest. R News, 2/3.

Maina, J., McClanahan, T. R., Venus, V., Ateweberhan, M., & Madin, J. (2011). Global gradients of coral exposure to environmental stresses and implications for local management. PLoS One, 6(8), e23064. 10.1371/journal.pone.0023064

Martinez Arbizu, P. (2020). pairwiseAdonis: Pairwise multilevel comparison using adonis. In (Version R package version 0.4)

Miller, A. W., & Richardson, L. L. (2014). Emerging coral diseases: a temperature-driven process? Marine Ecology, 36(3), 278–291. 10.1111/maec.12142

Miller, J., Muller, E., Rogers, C., Waara, R., Atkinson, A., Whelan, K., Patterson, M., & Witcher, B. (2009). Coral disease following massive bleaching in 2005 causes 60% decline in coral cover on reefs in the US Virgin Islands. Coral Reefs, 28, 925–937. 10.1007/s00338-009-0531-7

Morris, L. A., Voolstra, C. R., Quigley, K. M., Bourne, D. G., & Bay, L. K. (2019). Nutrient Availability and Metabolism Affect the Stability of Coral-Symbiodiniaceae Symbioses. Trends Microbiol, 27(8), 678–689. 10.1016/j.tim.2019.03.004

Muller, E. M., Bartels, E., & Baums, I. B. (2018). Bleaching causes loss of disease resistance within the threatened coral species. Elife, 7. 10.7554/eLife.35066

Muller, E. M., Rogers, C. S., Spitzack, A. S., & van Woesik, R. (2008). Bleaching increases likelihood of disease on (Lamarck) in Hawksnest Bay, St John, US Virgin Islands. Coral Reefs, 27(1), 191–195. 10.1007/s00338-007-0310-2

Muller, E. M., Sartor, C., Alcaraz, N. I., & van Woesik, R. (2020). Spatial Epidemiology of the Stony-Coral-Tissue-Loss Disease in Florida. Frontiers in Marine Science, 7. 10.3389/fmars.2020.00163

Muller, E. M., & van Woesik, R. (2009). Shading reduces coral-disease progression. Coral Reefs, 28(3), 757–760. 10.1007/s00338-009-0504-x

Mydlarz, L. D., & Harvell, C. D. (2007). Peroxidase activity and inducibility in the sea fan coral exposed to a fungal pathogen. Comp Biochem Physiol A Mol Integr Physiol, 146(1), 54–62. 10.1016/j.cbpa.2006.09.005

Neely, K. L., Macaulay, K. A., Hower, E. K., & Dobler, M. A. (2020). Effectiveness of topical antibiotics in treating corals affected by Stony Coral Tissue Loss Disease. PeerJ, 8, e9289. 10.7717/peerj.9289

Oksanen, J., Simpson, G., Blanchet, F., Kindt, R., Legendre, P., Minchin, P., O’Hara, R., Solymos, P., Stevens, M., Szoecs, E., Wagner, H., Barbour, M., Bedward, M., Bolker, B., Borcard, D., Borman, T., Carvalho, G., Chirico, M., De Caceres, M., . . . Weedon, J. (2025). vegan: Community Ecology Package. In (Version R package version 2.8-0) https://vegandevs.github.io/vegan/

Page, C. E., Leggat, W., Egan, S., & Ainsworth, T. D. (2023). A coral disease outbreak highlights vulnerability of remote high-latitude lagoons to global and local stressors. iScience, 26(3), 106205. 10.1016/j.isci.2023.106205

Palacio-Castro, A. M., Soderberg, N., Zagon, Z., Cooke, K., Studivan, M. S., Gill, T., Kelble, C., Christian, T., & Enochs, I. C. (2025). Elevated temperature decreases stony coral tissue loss disease transmission, with little effect of nutrients. Scientific Reports, 15(1), 22261. 10.1038/s41598-025-06322-0

Palmer, C. V., Bythell, J. C., & Willis, B. L. (2010). Levels of immunity parameters underpin bleaching and disease susceptibility of reef corals. FASEB J, 24(6), 1935–1946. 10.1096/fj.09-152447

Palmer, C. V., Bythell, J. C., & Willis, B. L. (2011). A comparative study of phenoloxidase activity in diseased and bleached colonies of the coral Acropora millepora. Dev Comp Immunol, 35(10), 1098–1101. 10.1016/j.dci.2011.04.001

Palmer, C. V., McGinty, E. S., Cummings, D. J., Smith, S. M., Bartels, E., & Mydlarz, L. D. (2011). Patterns of coral ecological immunology: variation in the responses of Caribbean corals to elevated temperature and a pathogen elicitor. J Exp Biol, 214(Pt 24), 4240–4249. 10.1242/jeb.061267

Palmer, C. V., Mydlarz, L. D., & Willis, B. L. (2008). Evidence of an inflammatory-like response in non-normally pigmented tissues of two scleractinian corals. Proc Biol Sci, 275(1652), 2687–2693. 10.1098/rspb.2008.0335

Pinzon, C. J., Beach-Letendre, J., Weil, E., & Mydlarz, L. D. (2014). Relationship between phylogeny and immunity suggests older Caribbean coral lineages are more resistant to disease. PLoS One, 9(8), e104787. 10.1371/journal.pone.0104787

Pollock, F. J., Lamb, J. B., Field, S. N., Heron, S. F., Schaffelke, B., Shedrawi, G., Bourne, D. G., & Willis, B. L. (2014). Sediment and turbidity associated with offshore dredging increase coral disease prevalence on nearby reefs. PLoS One, 9(7), e102498. 10.1371/journal.pone.0102498

Precht, W. F., Gintert, B. E., Robbart, M. L., Fura, R., & van Woesik, R. (2016). Unprecedented Disease-Related Coral Mortality in Southeastern Florida. Sci Rep, 6, 31374. 10.1038/srep31374

Randall, C. J., & van Woesik, R. (2015). Contemporary white-band disease in Caribbean corals driven by climate change. Nature Climate Change, 5(4), 375–379. 10.1038/nclimate2530

Randazzo-Eisemann, A., Garza-Perez, J. R., & Figueroa-Zavala, B. (2022). The role of coral diseases in the flattening of a Caribbean Coral Reef over 23 years. Mar Pollut Bull, 181, 113855. 10.1016/j.marpolbul.2022.113855

Richardson, L. L. (1998). Coral diseases: what is really known? Trends Ecol Evol, 13(11), 438–443. 10.1016/s0169-5347(98)01460-8

Ristaino, J. B., Anderson, P. K., Bebber, D. P., Brauman, K. A., Cunniffe, N. J., Fedoroff, N. V., Finegold, C., Garrett, K. A., Gilligan, C. A., Jones, C. M., Martin, M. D., MacDonald, G. K., Neenan, P., Records, A., Schmale, D. G., Tateosian, L., & Wei, Q. (2021). The persistent threat of emerging plant disease pandemics to global food security. Proc Natl Acad Sci U S A, 118(23). 10.1073/pnas.2022239118

Rodenberg, C. A., Hajek, A. E., Nadel, H., Stefanski, A., Reich, P. B., & Haynes, K. J. (2024). Rising temperatures may increase fungal epizootics in northern populations of the invasive spongy moth in North America. NeoBiota, 95, 291–308. 10.3897/neobiota.95.126311

Roussin-Leveillee, C., Rossi, C. A. M., Castroverde, C. D. M., & Moffett, P. (2024). The plant disease triangle facing climate change: a molecular perspective. Trends Plant Sci, 29(8), 895–914. 10.1016/j.tplants.2024.03.004

Rowley, A. F., Baker-Austin, C., Boerlage, A. S., Caillon, C., Davies, C. E., Duperret, L., Martin, S. A. M., Mitta, G., Pernet, F., Pratoomyot, J., Shields, J. D., Shinn, A. P., Songsungthong, W., Srijuntongsiri, G., Sritunyalucksana, K., Vidal-Dupiol, J., Uren Webster, T. M., Taengchaiyaphum, S., Wongwaradechkul, R., & Coates, C. J. (2024). Diseases of marine fish and shellfish in an age of rapid climate change. iScience, 27(9), 110838. 10.1016/j.isci.2024.110838

Ruiz-Moreno, D., Willis, B. L., Page, A. C., Weil, E., Croquer, A., Vargas-Angel, B., Jordan-Garza, A. G., Jordan-Dahlgren, E., Raymundo, L., & Harvell, C. D. (2012). Global coral disease prevalence associated with sea temperature anomalies and local factors. Dis Aquat Organ, 100(3), 249–261. 10.3354/dao02488

Santos Ede, O., Alves, N., Jr., Dias, G. M., Mazotto, A. M., Vermelho, A., Vora, G. J., Wilson, B., Beltran, V. H., Bourne, D. G., Le Roux, F., & Thompson, F. L. (2011). Genomic and proteomic analyses of the coral pathogen Vibrio coralliilyticus reveal a diverse virulence repertoire. ISME J, 5(9), 1471–1483. 10.1038/ismej.2011.19

Sato, Y., Bourne, D. G., & Willis, B. L. (2009). Dynamics of seasonal outbreaks of black band disease in an assemblage of species at Pelorus Island (Great Barrier Reef, Australia). Proceedings of the Royal Society B-Biological Sciences, 276(1668), 2795–2803. 10.1098/rspb.2009.0481

Schneider, C. A., Rasband, W. S., & Eliceiri, K. W. (2012). NIH Image to ImageJ: 25 years of image analysis. Nat Methods, 9(7), 671–675. 10.1038/nmeth.2089

Sharoni, T., Jaimes-Becerra, A., Lewandowska, M., Aharoni, R., Voolstra, C. R., Fine, M., & Moran, Y. (2025). Heat Stress Drives Rapid Viral and Antiviral Innate Immunity Activation in Hexacorallia. Mol Ecol, 34(20), e70098. 10.1111/mec.70098

Shilling, E. N., Combs, I. R., & Voss, J. D. (2021). Assessing the effectiveness of two intervention methods for stony coral tissue loss disease on Montastraea cavernosa. Scientific Reports, 11(1), 8566. 10.1038/s41598-021-86926-4

Soderhall, K., & Cerenius, L. (1998). Role of the prophenoloxidase-activating system in invertebrate immunity. Curr Opin Immunol, 10(1), 23–28. 10.1016/s0952-7915(98)80026-5

Solórzano, L. (1969). Determination of ammonia in natural waters by the phenol-hypochlorite method. Limnology and Oceanography, 14, 799–801. 10.4319/lo.1969.14.5.0799

Takagi, T., Yoshioka, Y., Zayasu, Y., Satoh, N., & Shinzato, C. (2020). Transcriptome Analyses of Immune System Behaviors in Primary Polyp of Coral Acropora digitifera Exposed to the Bacterial Pathogen Vibrio coralliilyticus under Thermal Loading. Mar Biotechnol (NY), 22(6), 748–759. 10.1007/s10126-020-09984-1

Team, Q. D. (2025). *QGIS Geographic Information System*. In Open Source Geospatial Foundation Project. http://qgis.osgeo.org

Team, R. C. (2024). *R: A Language and Environment for Statistical Computing*. In R Foundation for Statistical Computing. https://www.R-project.org/

Traylor-Knowles, N., & Connelly, M. T. (2017). What Is Currently Known About the Effects of Climate Change on the Coral Immune Response. Current Climate Change Reports, 3(4), 252–260. 10.1007/s40641-017-0077-7

Ushijima, B., Gunasekera, S. P., Meyer, J. L., Tittl, J., Pitts, K. A., Thompson, S., Sneed, J. M., Ding, Y., Chen, M., Jay Houk, L., Aeby, G. S., Häse, C. C., & Paul, V. J. (2023). Chemical and genomic characterization of a potential probiotic treatment for stony coral tissue loss disease. Communications Biology, 6(1), 248. 10.1038/s42003-023-04590-y

van de Water, J., Chaib De Mares, M., Dixon, G. B., Raina, J. B., Willis, B. L., Bourne, D. G., & van Oppen, M. J. H. (2018). Antimicrobial and stress responses to increased temperature and bacterial pathogen challenge in the holobiont of a reef-building coral. Mol Ecol, 27(4), 1065–1080. 10.1111/mec.14489

van de Water, J. A., Lamb, J. B., van Oppen, M. J., Willis, B. L., & Bourne, D. G. (2015). Comparative immune responses of corals to stressors associated with offshore reef-based tourist platforms. Conserv Physiol, 3(1), cov032. 10.1093/conphys/cov032

van de Water, J. A. J. M., Lamb, J. B., Heron, S. F., van Oppen, M. J. H., & Willis, B. L. (2016). Temporal patterns in innate immunity parameters in reef-building corals and linkages with local climatic conditions. Ecosphere, 7(11). 10.1002/ecs2.1505

van Woesik, R., & Randall, C. J. (2017). Coral disease hotspots in the Caribbean. Ecosphere, 8(5). 10.1002/ecs2.1814

Vega Thurber, R., Mydlarz, L. D., Brandt, M., Harvell, D., Weil, E., Raymundo, L., Willis, B. L., Langevin, S., Tracy, A. M., Littman, R., Kemp, K. M., Dawkins, P., Prager, K. C., Garren, M., & Lamb, J. (2020). Deciphering Coral Disease Dynamics: Integrating Host, Microbiome, and the Changing Environment. Frontiers in Ecology and Evolution, 8. 10.3389/fevo.2020.575927

Vega Thurber, R. L., Burkepile, D. E., Fuchs, C., Shantz, A. A., McMinds, R., & Zaneveld, J. R. (2014). Chronic nutrient enrichment increases prevalence and severity of coral disease and bleaching. Global Change Biology, 20(2), 544–554. 10.1111/gcb.12450

Voss, J. D., & Richardson, L. L. (2006). Nutrient enrichment enhances black band disease progression in corals. Coral Reefs, 25(4), 569–576. 10.1007/s00338-006-0131-8

Walker, B. K., Turner, N. R., Noren, H. K. G., Buckley, S. F., & Pitts, K. A. (2021). Optimizing Stony Coral Tissue Loss Disease (SCTLD) Intervention Treatments on Montastraea cavernosa in an Endemic Zone. Frontiers in Marine Science, 8. 10.3389/fmars.2021.666224

Walton, C. J., Hayes, N. K., & Gilliam, D. S. (2018). Impacts of a Regional, Multi-Year, Multi-Species Coral Disease Outbreak in Southeast Florida. Frontiers in Marine Science, 5. 10.3389/fmars.2018.00323

Watch, N. C. R. (2019). NOAA Coral Reef Watch Version 3.1 Daily 5km Satellite Regional Virtual Station Time Series Data for Southeast Florida, Mar. 12, 2013-Mar. 11, 2014 NOAA Coral Reef Watch. https://coralreefwatch.noaa.gov/product/vs/data.php

Wear, S. L., & Thurber, R. V. (2015). Sewage pollution: mitigation is key for coral reef stewardship. Ann N Y Acad Sci, 1355(1), 15–30. 10.1111/nyas.12785

Weil, E. (2004). Coral Reef Diseases in the Wider Caribbean. In E. Rosenberg & Y. Loya (Eds.), Coral Health and Disease (pp. 35–68). Springer Berlin Heidelberg. 10.1007/978-3-662-06414-6_2

Wickham, H. (2016). *ggplot2: Elegant Graphics for Data Analysis*. Springer-Verlag.

Williams, S. D., Walter, C. S., & Muller, E. M. (2021). Fine Scale Temporal and Spatial Dynamics of the Stony Coral Tissue Loss Disease Outbreak Within the Lower Florida Keys. Frontiers in Marine Science, 8. 10.3389/fmars.2021.631776

