## Supplementary material for "Environmental conditions modulate constitutive and induced host immunity contributing to spatial patterns of coral disease susceptibility": Table S

**Table S1.** Environmental summary of predictors used in disease severity full random forest model. Variable = type of environmental variable, predictor = metric of that variable, time period = temporal duration predictor calculated over. e.g., Chlorophyll-a maximum year prior, is the maximum Chlorophyll-a concentration within the year prior to the initial benthic survey used to count *Montastraea cavernosa* abundance. PAR maximum during, is the maximum PAR measurement in the two years between monitoring surveys.

| Variable | Predictor | Time period | Mean | SD |
|---|---|---|---|---|
| Chlorophyll-a (mg $m^{-3}$) | Maximum | Year prior | 8.88 | 10.95 |
| PAR (Einstein $m^{-2}$ $day^{-1}$) | Maximum | During | 59.09 | 1.32 |
| PAR (Einstein $m^{-2}$ $day^{-1}$) | Mean | During | 43.52 | 1.55 |
| Temperature (°C) | Maximum | Year prior | 31.07 | 0.9 |
| Temperature (°C) | Maximum | During | 31.37 | 0.71 |
| Temperature (°C) | Mean | Year prior | 26.97 | 0.94 |
| Temperature (°C) | Mean | During | 27.07 | 0.35 |
| Temperature (°C) | Minimum | Year prior | 21.89 | 1.93 |
| Temperature (°C) | Minimum | During | 21.24 | 1.3 |
| Thermal stress (days) | Heat stress | Year prior | 4.03 | 7.41 |
| Thermal stress (days) | Heat stress | During | 6.68 | 11.89 |
| Thermal stress (days) | Cold stress | Year prior | 2.21 | 6.15 |
| Thermal stress (days) | Cold stress | During | 4.39 | 10.36 |
| Thermal stress (days) | Maximum DHW | During | 1.73 | 2.15 |

**Table S2.** List of environmental treatment types including final temperature and ammonia values. Each treatment type corresponds to a single experimental tank. Treatments listed were crossed with immune challenge treatment (placebo or immune challenge) in a full factorial manner for a final total of 12 treatment groups.

| Treatment | Temperature | Ammonia |
|---|---|---|
| Ambient, None | 28 °C | 0 |
| Ambient, Mid | 28 °C | 0.01 mg/L |
| Ambient, High | 28 °C | 0.05 mg/L |
| Heat, None | 32 °C | 0 |
| Heat, Mid | 32 °C | 0.01 mg/L |
| Heat, None | 32 °C | 0.05 mg/L |

**Table S3.** LMM results for the impacts of temperature, nutrients, and their interaction on symbiont density. DF = degrees of freedom; Num = Numerator, Den = denominator. Bold font indicates significant p-values. *p < 0.05, **p < 0.001.

| **Factor** | **Sum of Squares** | **Mean Squares** | **NumDF** | **DenDF** | **F value** | **p-value** |
|---|---|---|---|---|---|---|
| Temperature | 7583436 | 7583436 | 1 | 72 | 40.99 | **<0.001**** |
| Nutrients | 215628 | 107814 | 2 | 72 | 0.58 | 0.56 |
| Temperature*Nutrients | 1057430 | 528715 | 2 | 72 | 2.87 | 0.06 |

**Table S4.** PERMANOVA results for combined immunological metrics. Df = degrees of freedom. $R^2$ value gives an estimate of the amount of variance explained by a given factor. Bold font indicates significant p-values. *p < 0.05, **p < 0.001.

| **Factor** | **Df** | **Sum of Squares** | **$R^2$** | **F value** | **p-value** |
|---|---|---|---|---|---|
| Temperature | 1 | 52.55 | 0.11 | 10.15 | **0.001**** |
| Nutrients | 2 | 21.04 | 0.042 | 2.03 | **0.03*** |
| Infection | 1 | 4.14 | 0.008 | 0.80 | 0.54 |
| Temp*Nutrients | 2 | 23.05 | 0.046 | 2.23 | **0.01*** |
| Temp*Infection | 1 | 3.87 | 0.008 | 0.75 | 0.61 |
| Nutrients*Immune Challenge | 2 | 17.90 | 0.036 | 1.73 | 0.06 |
| Temp*Nutrients*Immune Challenge | 2 | 2.72 | 0.006 | 0.26 | 0.99 |
| Residual | 72 | 372.7 | 0.75 | | |
| Total | 83 | 498.0 | 1.00 | | |

**Table.** Post-hoc analysis of PERMANOVA results for the interaction of Temp*Nutrients. Results are shown for all pairwise comparisons. Bold font indicates significant p-values. *p < 0.05, **p < 0.001.

| | Ambient, None | Ambient, Mid | Ambient, High | Heat, None | Heat,Mid |
|---|---|---|---|---|---|
| Ambient, Mid | 1.00 | | | | |
| Ambient, High | 1.00 | 0.585 | | | |
| Heat, None | 0.255 | **0.045*** | **0.045*** | | |
| Heat, Mid | **0.030*** | **0.015*** | **0.015*** | 1.00 | |
| Heat,High | 0.105 | **0.015*** | **0.015*** | 1.00 | **0.030*** |

**Table S6.** PCA results showing standard deviation and proportion of variance explained by each PC

| | **PC1** | **PC2** | **PC3** | **PC4** | **PC5** | **PC6** |
|---|---|---|---|---|---|---|
| **General Statistics** | | | | | | |
| Standard Deviation | 1.30 | 1.12 | 1.05 | 0.88 | 0.85 | 0.68 |
| Proportion of Variance | 0.28 | 0.21 | 0.18 | 0.13 | 0.12 | 0.08 |
| Cumulative Proportion | 0.28 | 0.49 | 0.67 | 0.80 | 0.92 | 1.00 |
| **Loadings (eigenvectors)** | | | | | | |
| Symbiont Density | 0.45 | 0.56 | 0.06 | -0.34 | -0.03 | 0.60 |
| Catalase | -0.25 | 0.46 | 0.58 | -0.06 | 0.54 | -0.30 |
| Peroxidase | 0.58 | -0.16 | 0.17 | -0.49 | -0.15 | -0.58 |
| Total Phenoloxidase | 0.31 | -0.13 | 0.66 | 0.57 | -0.32 | 0.14 |
| Melanin | 0.28 | 0.34 | -0.41 | 0.51 | -0.08 | -0.41 |
| Antibacterial Activity | 0.47 | 0.56 | -0.14 | 0.23 | 0.76 | 0.14 |

**Table S7.** LMM results investigating variance in each principal component as a result of temperature, nutrients, immune challenge and their interactions. DF = degrees of freedom; Num = Numerator, Den = denominator. Bold font indicates significant p-value. *p < 0.05, **p < 0.001.

| **PC1** | | | | | | |
|---|---|---|---|---|---|---|
| **Effect** | **Sum of Squares** | **Mean Squares** | **NumDF** | **DenDF** | **F value** | **p-value** |
| Temperature | 48.95 | 48.95 | 1 | 66 | 56.44 | **<0.001**** |
| Nutrients | 1.04 | 0.520 | 2 | 66 | 0.599 | 0.552 |
| Infection | 0.04 | 0.039 | 1 | 66 | 0.445 | 0.834 |
| Temperature: Nutrients | 1.34 | 0.670 | 2 | 66 | 0.772 | 0.466 |
| Temperature: Infection | 0.01 | 0.013 | 1 | 66 | 0.016 | 0.901 |
| Nutrients: Infection | 2.16 | 1.080 | 2 | 66 | 1.246 | 0.294 |
| Temperature: Nutrients: Infection | 0.23 | 0.114 | 2 | 66 | 0.131 | 0.877 |
| **PC2** | | | | | | |
| **Effect** | **Sum of Squares** | **Mean Squares** | **NumDF** | **DenDF** | **F value** | **p-value** |
| Temperature | 0.711 | 0.711 | 1 | 66 | 0.792 | 0.377 |
| Nutrients | 1.080 | 0.540 | 2 | 66 | 0.601 | 0.551 |
| Infection | 0.819 | 0.819 | 1 | 66 | 0.912 | 0.343 |
| Temperature: Nutrients | 12.50 | 6.25 | 2 | 66 | 6.958 | **0.002*** |
| Temperature: Infection | 0.441 | 0.441 | 1 | 66 | 0.491 | 0.486 |
| Nutrients: Infection | 3.25 | 1.62 | 2 | 66 | 1.808 | 0.172 |
| Temperature: Nutrients: Infection | 0.869 | 0.434 | 2 | 66 | 0.483 | 0.619 |
| **PC3** | | | | | | |
| **Effect** | **Sum of Squares** | **Mean Squares** | **NumDF** | **DenDF** | **F value** | **p-value** |
| Temperature | 0.750 | 0.750 | 1 | 66 | 0.865 | 0.356 |
| Nutrients | 16.09 | 8.04 | 2 | 66 | 9.280 | **<0.001**** |
| Infection | 0.767 | 0.767 | 1 | 66 | 0.885 | 0.350 |
| Temperature: Nutrients | 0.687 | 0.344 | 2 | 66 | 0.396 | 0.674 |
| Temperature: Infection | 0.721 | 0.721 | 1 | 66 | 0.831 | 0.365 |
| Nutrients: Infection | 5.237 | 2.618 | 2 | 66 | 3.021 | 0.556 |
| Temperature: Nutrients: Infection | 0.224 | 0.112 | 2 | 66 | 0.129 | 0.879 |
| **PC4** | | | | | | |
| **Effect** | **Sum of Squares** | **Mean Squares** | **NumDF** | **DenDF** | **F value** | **p-value** |
| Temperature | 1.596 | 1.596 | 1 | 66 | 2.619 | 0.110 |
| Nutrients | 1.447 | 0.723 | 2 | 66 | 1.187 | 0.312 |
| Infection | 0.304 | 0.304 | 1 | 66 | 0.499 | 0.482 |
| Temperature: Nutrients | 7.134 | 3.567 | 2 | 66 | 5.852 | **0.005*** |
| Temperature: Infection | 0.542 | 0.542 | 1 | 66 | 0.889 | 0.349 |
| Nutrients: Infection | 5.765 | 2.882 | 2 | 66 | 4.729 | **0.012*** |
| Temperature: Nutrients: Infection | 0.390 | 0.195 | 2 | 66 | 0.320 | 0.727 |

**Table S8.** Results of post-hoc testing for significant model terms from LMMs of the impacts of temperature, nutrients, immune challenge and their interactions on PC scores. Df = degrees of freedom. Bold font indicates significant p-value. *p < 0.05, **p < 0.001.

| **PC2- Temperature:Nutrients** | | | | | |
|---|---|---|---|---|---|
| **Pairwise comparison** | **Estimate** | **SE** | **Df** | **t ratio** | **p-value** |
| None Ambient - None Heat | -0.387 | 0.358 | 66 | -1.08 | 0.888 |
| None Ambient - Mid Heat | -0.616 | 0.358 | 66 | -1.72 | 0.524 |
| None Ambient - High Heat | 0.412 | 0.358 | 66 | 1.15 | 0.858 |
| Mid Ambient - None Ambient | 0.280 | 0.358 | 66 | 0.782 | 0.970 |
| Mid Ambient - None Heat | -0.106 | 0.358 | 66 | -0.297 | 1.000 |
| Mid Ambient - Mid Heat | -0.336 | 0.358 | 66 | -0.938 | 0.935 |
| Mid Ambient - High Heat | 0.692 | 0.358 | 66 | 1.933 | 0.392 |
| Mid Heat - None Heat | 0.229 | 0.358 | 66 | 0.641 | 0.987 |
| High Ambient - None Ambient | 0.863 | 0.358 | 66 | 2.408 | 0.169 |
| High Ambient - Mid Ambient | 0.582 | 0.358 | 66 | 1.626 | 0.585 |
| High Ambient - None Heat | 0.476 | 0.358 | 66 | 1.329 | 0.768 |
| High Ambient - Mid Heat | 0.246 | 0.358 | 66 | 0.688 | 0.983 |
| High Ambient - High Heat | 1.275 | 0.358 | 66 | 3.558 | **0.009*** |
| High Heat - None Heat | -0.799 | 0.358 | 66 | -2.23 | 0.238 |
| High Heat - Mid Heat | -1.028 | 0.358 | 66 | -2.87 | 0.059 |
| **PC3- Nutrients** | | | | | |
| **Pairwise comparison** | **Estimate** | **SE** | **Df** | **t ratio** | **p-value** |
| Mid-None | 0.593 | 0.249 | 66 | 2.384 | 0.052 |
| High-None | -0.477 | 0.249 | 66 | -1.915 | 0.142 |
| High-Mid | -1.070 | 0.249 | 66 | -4.300 | **<0.001**** |
| **PC4- Temperature:Nutrients** | | | | | |
| **Pairwise comparison** | **Estimate** | **SE** | **Df** | **t ratio** | **p-value** |
| None Ambient - None Heat | -0.098 | 0.295 | 66 | -0.333 | 0.999 |
| None Ambient - Mid Heat | -0.513 | 0.295 | 66 | -1.74 | 0.511 |
| None Ambient - High Heat | -0.122 | 0.295 | 66 | -0.413 | 0.998 |
| Mid Ambient - None Ambient | -0.548 | 0.295 | 66 | -1.858 | 0.437 |
| Mid Ambient - None Heat | -0.646 | 0.295 | 66 | -2.191 | 0.256 |
| Mid Ambient - Mid Heat | -1.062 | 0.295 | 66 | -3.597 | **0.008*** |
| Mid Ambient - High Heat | -0.670 | 0.295 | 66 | -2.271 | 0.221 |
| Mid Heat - None Heat | 0.415 | 0.295 | 66 | 1.407 | 0.723 |
| High Ambient - None Ambient | 0.455 | 0.295 | 66 | 1.54 | 0.640 |
| High Ambient - Mid Ambient | 1.003 | 0.295 | 66 | 3.398 | **0.014*** |
| High Ambient - None Heat | 0.356 | 0.295 | 66 | 1.207 | 0.832 |
| High Ambient - Mid Heat | -0.059 | 0.295 | 66 | -0.199 | 1.000 |
| High Ambient - High Heat | 0.333 | 0.295 | 66 | 1.128 | 0.868 |
| High Heat - None Heat | 0.024 | 0.295 | 66 | 0.08 | 1.000 |
| High Heat - Mid Heat | -0.392 | 0.295 | 66 | -1.327 | 0.769 |
| **PC4- Nutrients:Infection** | | | | | |

| Pairwise comparison | Estimate | SE | Df | t ratio | p-value |
|---|---|---|---|---|---|
| Mid Control - None Control | 0.039 | 0.295 | 66 | 0.133 | 1.00 |
| High Control - None Control | -0.256 | 0.295 | 66 | -0.868 | 0.953 |
| High Control - Mid Control | -0.295 | 0.295 | 66 | -1.001 | 0.916 |
| None Bacteria - None Control | -0.139 | 0.295 | 66 | -0.472 | 0.997 |
| None Bacteria - Mid Control | -0.179 | 0.295 | 66 | -0.605 | 0.990 |
| None Bacteria - High Control | 0.117 | 0.295 | 66 | 0.396 | 0.999 |
| Mid Bacteria - None Control | -0.312 | 0.295 | 66 | -1.056 | 0.897 |
| Mid Bacteria - None Bacteria | -0.172 | 0.295 | 66 | -0.584 | 0.992 |
| Mid Bacteria - Mid Control | -0.351 | 0.295 | 66 | -1.189 | 0.841 |
| Mid Bacteria - High Control | -0.055 | 0.295 | 66 | -0.188 | 1.00 |
| High Bacteria - None Bacteria | 0.734 | 0.295 | 66 | 2.489 | 0.142 |
| High Bacteria - Mid Bacteria | 0.907 | 0.295 | 66 | 3.072 | **0.035*** |
| High Bacteria - None Control | 0.595 | 0.295 | 66 | 2.016 | 0.344 |
| High Bacteria - Mid Control | 0.556 | 0.295 | 66 | 1.884 | 0.421 |
| High Bacteria - High Control | 0.851 | 0.295 | 66 | 2.884 | 0.057 |

**Table S9.** Results of post-hoc testing for significant model terms from LMMs of the impacts of temperature, nutrients, immune challenge and their interactions on individual immune metrics. Df = degrees of freedom. Bold font indicates significant p-value. *p < 0.05, **p < 0.001.

| **Catalase- Nutrients** | | | | | |
|---|---|---|---|---|---|
| **Pairwise comparison** | **Estimate** | **SE** | **Df** | **t ratio** | **p-value** |
| Mid-None | 0.184 | 0.225 | 66 | 0.816 | 0.694 |
| High-None | -0.526 | 0.225 | 66 | -2.340 | 0.057 |
| High-Mid | -0.710 | 0.225 | 66 | -3.157 | **0.007*** |
| **Catalase- Nutrients:Infection** | | | | | |
| **Pairwise comparison** | **Estimate** | **SE** | **Df** | **t ratio** | **p-value** |
| Mid Control - None Control | -0.509 | 0.318 | 66 | -1.6 | 0.601 |
| High Control - None Control | -0.642 | 0.318 | 66 | -2.02 | 0.342 |
| High Control - Mid Control | -0.134 | 0.318 | 66 | -0.42 | 0.998 |
| None Bacteria - None Control | -0.607 | 0.318 | 66 | -1.907 | 0.407 |
| None Bacteria - Mid Control | -0.098 | 0.318 | 66 | -0.307 | 1.00 |
| None Bacteria - High Control | 0.036 | 0.318 | 66 | 0.113 | 1.00 |
| Mid Bacteria - None Control | 0.269 | 0.318 | 66 | 0.847 | 0.957 |
| Mid Bacteria - None Bacteria | 0.876 | 0.318 | 66 | 2.754 | 0.078 |
| Mid Bacteria - Mid Control | 0.778 | 0.318 | 66 | 2.447 | 0.155 |
| Mid Bacteria - High Control | 0.912 | 0.318 | 66 | 2.867 | 0.059 |
| High Bacteria - None Control | -1.017 | 0.318 | 66 | -3.197 | **0.025*** |
| High Bacteria - None Bacteria | -0.410 | 0.318 | 66 | -1.29 | 0.790 |
| High Bacteria - Mid Control | -0.508 | 0.318 | 66 | -1.597 | 0.604 |
| High Bacteria - Mid Bacteria | -1.286 | 0.318 | 66 | -4.044 | **0.002*** |
| High Bacteria - High Control | -0.374 | 0.318 | 66 | -1.177 | 0.846 |
| **Melanin- Temperature:Nutrients** | | | | | |
| **Pairwise comparison** | **Estimate** | **SE** | **Df** | **t ratio** | **p-value** |
| None Ambient - None Heat | -0.195 | 0.217 | 66 | -0.898 | 0.946 |
| None Ambient - Mid Heat | -0.186 | 0.217 | 66 | -0.857 | 0.955 |
| None Ambient - High Heat | -0.034 | 0.217 | 66 | -0.156 | 1.000 |
| Mid Ambient - None Ambient | -0.115 | 0.217 | 66 | -0.528 | 0.995 |
| Mid Ambient - None Heat | -0.310 | 0.217 | 66 | -1.425 | 0.712 |
| Mid Ambient - Mid Heat | -0.301 | 0.217 | 66 | -1.385 | 0.736 |
| Mid Ambient - High Heat | -0.149 | 0.217 | 66 | -0.684 | 0.983 |
| Mid Heat - None Heat | -0.009 | 0.217 | 66 | -0.041 | 1.000 |
| High Ambient - None Ambient | 0.670 | 0.217 | 66 | 3.083 | **0.034*** |
| High Ambient - Mid Ambient | 0.785 | 0.217 | 66 | 3.611 | **0.008*** |
| High Ambient - None Heat | 0.475 | 0.217 | 66 | 2.186 | 0.258 |
| High Ambient - Mid Heat | 0.484 | 0.217 | 66 | 2.226 | 0.240 |
| High Ambient - High Heat | 0.636 | 0.217 | 66 | 2.927 | 0.051 |
| High Heat - None Heat | -0.161 | 0.217 | 66 | -0.741 | 0.976 |
| High Heat - Mid Heat | -0.152 | 0.217 | 66 | -0.701 | 0.981 |
